# Early Detection of an Invasive Alien Plant (*Phragmites australis*) Using Unoccupied Aerial Vehicles and Artificial Intelligence

**DOI:** 10.1101/2024.06.06.593339

**Authors:** Antoine Caron-Guay, Mickaël Germain, Etienne Laliberté

## Abstract

The combination of unoccupied aerial vehicles (UAVs) and artificial intelligence to map vegetation represents a promising new approach to improve the detection of invasive alien plant species (IAPS). The high spatial resolution achievable with UAVs and recent innovations in computer vision, especially with convolutional neural networks, suggest that early detection of IAPS could be possible, thus facilitating their management. In this study, we evaluated the suitability of this approach for mapping the location of common reed (*Phragmites australis* subsp. *australis*) within a national park located in southern Quebec, Canada. We collected data on six distinct dates during the growing season, covering environments with different levels of reed invasion. Overall, model performance was high for the different dates and zones, especially for recall (mean of 0.89). The results showed an increase in performance, reaching a peak following the appearance of the inflorescence in September (highest F1-score at 0.98). Furthermore, a decrease in spatial resolution negatively affected recall (18% decrease between a spatial resolution of 0.15 cm pixel^−1^ and 1.50 cm pixel^−1^) but did not have a strong impact on precision (2% decrease). Despite challenges associated with common reed mapping in a post-treatment monitoring context, the use of UAVs and deep learning shows great potential for IAPS detection when supported by a suitable dataset. Our results show that, from an operational point of view, this approach could be an effective tool for speeding up the work of biologists in the field and ensuring better management of IAPS.

## 1. Introduction

Nearly 25% of plant and animal species are threatened with extinction, with many researchers describing a sixth mass extinction as currently underway (Barnosky et al., 2011; Ceballos et al., 2015; IPBES, 2019; Payne et al., 2016). Invasive alien species are second only to habitat loss as a cause of this biodiversity loss (Baillie et al., 2004; Bellard et al., 2016). Besides altering plant and animal communities structure, invasive alien plant species (IAPS) can have several environmental consequences such as disrupting ecosystem functions, disturbances regimes and biogeochemical cycles (Ehrenfeld, 2010; IPBES, 2023; Levine et al., 2003). There are also economic impacts, with global costs for invasive alien species estimated at US$34 billion per year for their management alone, which represents only 8% of the total estimated costs of $423 billion (IPBES, 2023). Even societal impacts can be considered with some IAPS changing the visual landscape and preventing outdoor activities (Pejchar & Mooney, 2009).

Since IAPS are often well-established in their new environment, management mainly involves constant monitoring to control them and rapidly eradicate new growth. However, current methods rely primarily on field surveys, with on-site visual observers, and potential distribution maps analysis (Gormley et al., 2011; Kattenborn, Lopatin, et al., 2019). These methods do not allow the survey of a large area at a fine scale, can take a considerable amount of time depending on the surface area to be covered, and can cost a great deal of money if monitoring is to be carried out over several years (Drake et al., 2003; Xie et al., 2008). With new technologies developing quickly, it is possible to design new approaches allowing early detection, rapid treatment, and long-term monitoring to ensure effective management of the target species.

Remote sensing is a technology that has been used for many years to monitor plant communities without requiring physical presence on the field, but the advent of unoccupied aerial vehicles (UAVs) that are readily available to the public have the potential to transform vegetation monitoring. The use of satellites and aircrafts allows data to be acquired at different spectral, spatial and temporal resolutions to map functional traits, biomes distribution, presence of species, or simply vegetation indexes (Cavender-Bares et al., 2022; Huylenbroeck et al., 2020; Xie et al., 2008). In the case of IAPS, aerial satellite and airborne imagery can be used to detect large colonies and estimate the degree of potential invasiveness over time to prevent spread (Elmer et al., 2021; Huang & Asner, 2009). However, these techniques may be limited in temporal and spatial resolutions (Matese et al., 2015; Mumby et al., 1999; S. Wang et al., 2023). In contrast, UAVs can capture imagery at very high spatial resolutions, reaching subcentimeter level. This facilitates species identification across various growth stages and helps differentiation between closely resembling species, at the individual level (Ding et al., 2023; James & Bradshaw, 2020). Additionally, the utilization of UAVs facilitates the establishment of a reproducible workflow for long-term monitoring endeavors. They offer high operational capability at a relatively low cost, allowing for the collection of data from challenging-to-reach locations (Sun et al., 2021). Together, these attributes suggest the possibility of early IAPS detection with UAVs. This early detection is even more crucial in the case of an invasive species that can spread exponentially, allowing rapid, targeted intervention before the population gets so large that eradication becomes very difficult (Bradley, 2014; Reaser et al., 2020).

One of the major challenges with remote sensing is the large amount of data generated that needs to be processed before results can be obtained. This is where artificial intelligence (AI), and computer vision, can assist in analyzing images by deriving descriptive information from visual data (Prince, 2012). Convolutional neural networks (CNNs) in particular have shown promise in this field (Brodrick et al., 2019; Katal et al., 2022; Kattenborn et al., 2021; Zhang et al., 2016). CNNs enable the extraction of complex features from images, such as color, texture, edge and shape of objects, through the use of filters and model layers (Kattenborn et al., 2021; Maggiori et al., 2017). These various low-level features are used to help the model in determining the optimal output target description. Furthermore, the ability of CNNs to analyze spatial patterns enhances its efficiency in processing high spatial resolution from UAVs data.

Considering phenological changes through time could also improve results with CNNs (Bradley, 2014). IAPS often have the ability to utilize environmental resources in a different way than native species, allowing them to establish themselves as dominant plants (Herbold & Moyle, 1986; Shea & Chesson, 2002). These differences between invasive and native species can be observed when considering temporal variations over the growing season. For example, IAPS can have a longer growing season, with an earlier start or a later end, or a shift in the period of flowering (Vymazal & Krőpfelová, 2005; Wolkovich & Cleland, 2011). Once the best moment for detection during the growing season has been found, the fully trained model can be seamlessly applied to new, unseen data (Kattenborn, Eichel, et al., 2019). The advantage of CNNs becomes particularly evident in this context by allowing the automation of monitoring over years without the need for additional annotations.

In this study, we focused on one IAPS in particular, the common reed (*Phragmites australis* (Cavanilles) Trinius ex Steudel subsp. *australis*). The main objective was to find out whether early detection of the species is possible, using an UAV to collect the data and artificial intelligence to process it. Imagery acquisition was carried out several times during the growing season to determine whether phenology could impact results and whether a particular time during the season was better for the detection. In addition, to assess the operational potential of this approach, a resampling of the ground sample distance (GSD) was carried out to find the best compromise between flight height and mission duration. We expected that specific features useful to identify the species in relation to other similar species in the field would improve model performance (Müllerová, Brůna, et al., 2017). Useful features such as inflorescences appear later in the season, and so we predicted that the best results would be obtained later in the summer, towards the end of the growing season. Furthermore, a decrease in spatial resolution should correspond to an equivalent decrease in model performance. The best results should be obtained with the highest resolution, particularly in the case of early detection, where higher resolution imagery is required (Bradley, 2014).

Several studies have already demonstrated that the use of UAVs and AI can effectively identify IAPS (James & Bradshaw, 2020; Kattenborn, Lopatin, et al., 2019). Other studies focusing on this approach specifically in the case of *Phragmites* detection have reached the same conclusion (Anderson et al., 2021; Cohen & Lewis, 2020; Higgisson et al., 2021; Mohler & Morse, 2022; Samiappan et al., 2017). Most studies, however, do not use very high-resolution imagery (subcentimer level), the highest resolution being 3 cm pixel^−1^ (Anderson et al., 2021; Higgisson et al., 2021). Moreover, no study has verified detection in the context of a management zone, where results may not be as good. In these previous studies, detection was carried out mainly on the reed colonies level, while the species is primarily found as isolated stems or regrowth that has resisted treatment in management zones where annual treatment against the species is conducted. Overall, the purpose of our study was to determine whether this approach is suitable from an operational point of view as a replacement for conventional field surveys for common reed management.

## 2. Methods

### 2.1 Study site

This study took place at the Parc national des Îles-de-Boucherville, a protected area of approximately 8 km^2^ made up of five islands in the Saint Lawrence River in the Montréal area, Québec, Canada (45°35’14”N 73°28’58”W). This national park was created in 1984, but much of the island was actively farmed at the time, and agricultural activities were gradually abandoned. Over the years, these abandoned fields underwent secondary succession and became dominated by herbs and shrubs, with a small number of scattered trees inland (Pellerin et al., 2016). There are also few young woodlands within the area and a single mature woodland on Île Grosbois with trees aged between 70 and 90 years (Laliberté et al., 2006; Ross, 1990). In the past few years, the park has begun restoring most of the remaining farmland to natural conditions (Karathanos et al., 2015; Sépaq, 2021). However, each time an agricultural field is abandoned, the bare ground creates ideal conditions for the establishment of exotic invasive plant species. Nowadays, the presence of several exotic invasive plant species has been documented within the park, such as garlic mustard, Japanese knotweed, European buckthorn, glossy buckthorn as well as common reed across the islands (Sépaq, 2022). Of all these exotic species, common reed is of primary concern to the park authorities and is subject to a control program. Other species found include goldenrod, common milkweed and reed canary grass, the latter can also be considered exotic because of its non-native genotype (Lavergne & Molofsky, 2004).

### 2.2 Target invasive alien plant

Common reed is a cosmopolitan species of the Poaceae family. Growing up to 5 m tall, it will often dominate the herbaceous stratum, forming large and dense monospecific colonies (Mal & Narine, 2004; Packer et al., 2017). While scientists report a dieback of the species in its native distribution in Europe, it is considered as an invasive alien species in North America, where it continues to spread within the ecosystems (Canadian Food Inspection Agency, 2008; Gigante et al., 2011). The first observations of the exotic genotype in Québec dates back to the early 20th century, but its rapid dispersal started in the 1970s, mainly with the development of the road network (Jodoin et al., 2008; Lelong et al., 2007). Seed dispersal and the accidental transport of stem or rhizome fragments, which can in turn produce a new clone, are the main methods by which the species establishes itself in a new environment (Brisson et al., 2008). Once established, the species can rapidly spread and invade a large part of the ecosystem, via stolons, also called “legehalme”, and rhizomes that link the emerging stems to the colony (Clevering & Van Der Toorn, 2000; Maheu-Giroux & Blois, 2005; Mal & Narine, 2004). Its rhizome is very dense, making up more than half of its biomass, which favours its vegetative growth and hinders its eradication (Lavoie et al., 2019; Packer et al., 2017). Whereas the native subspecies, *Phragmites australis* subsp. *americanus* Saltonstall, P.M. Peterson & Soreng, accounted for almost 90% of colonies of reed in Québec before 1950, it is now estimated that it represents less than 5% (Lelong et al., 2007). For the scope of this study, American reed is not present in the national park and its potential impact on the detection model for the exotic subspecies will therefore not be considered.

The common reed now occupies a large part of the park’s territory, growing from 1 ha in 1980 to 33 ha in 2002 and 86 ha in 2010, with the Îles-de-Boucherville considered the largest reed bed in the province (Hudon et al., 2005; Lavoie et al., 2019; Tougas-Tellier, 2013). An annual monitoring and treatment are carried out in accordance with the park’s management strategy against this species in certain areas to reduce its spread. To achieve this, a team of biologists surveys these areas for new shoots of common reed and then eradicates them using glyphosate applied by hand on the stems and foliage. Lastly, other grasses present in the study area, such as redtop, bluejoint reedgrass and reed canary grass, may be confused with the target species. At the vegetative stage, the broad leaves of common reed and its hairy ligule are generally distinguishing features, while the bushy, most often gray or purple-coloured panicle that appears at flowering is a distinctive feature that helps to make the differentiation quickly (Mal & Narine, 2004).

### 2.3 Imagery acquisition and processing

Imagery was acquired using a DJI Matrice 300 RTK UAV equipped with a DJI Zenmuse P1 camera (DJI Science and Technology Co. Ltd., Shenzhen, China). The P1 consists of a 45 MP sensor with a 35 mm lens producing RGB images of 8192 by 5460 pixels (px). Flight missions were carried out from July to October 2022 on six dates and at a height of 12 m above ground level. This altitude was selected to obtain the highest possible spatial resolution (approximately 1.5 mm px^−1^) while minimizing the impact on vegetation of wind generated by the drone’s propellers. It is also the lowest altitude possible when planning a mission using DJI Pilot 2 on the remote controller. The values for frontal and lateral overlap during missions were 90% and 80% respectively. Three distinct zones, named ‘Dense established’, ‘Establishing’ and ‘Post-treatment’ thereafter, were chosen to represent the variability of reed presence in the park (Supplementary Fig. S8). They are all former agricultural fields that were abandoned in 2004, 2008 and 2012 respectively.

’Dense established’ is a zone where the species is well established, forming colonies with a high density of stems. Diversity is rather low in this area and there are no shrubs. ‘Establishing’ is a site where the presence of reed is highly variable. Few large colonies are present in the middle of the field and in drainage ditches, while the presence of the species elsewhere in the field is sporadic. There is also a relatively high plant species and structural diversity at this location, with presence of different herbs, shrubs, and small trees. Finally, ‘Post-treatment’ is a 40 ha area where the common reed management strategy is applied. Because of this treatment, the presence of the species is mainly in the form of sparse young shoots across the field that have regrown from treated colonies, while other grass species are more dominant.

Within each zone, 4, 20 and 5 missions respectively were carried out over areas ranging from 50 to 600 m^2^, with an average surface area of 225 m^2^ per mission (Supplementary Figure S1). The missions were planned to cover as much of the variability of *Phragmites* colonization as possible. Ground control points (GCPs) were added for some of the missions to verify the accuracy of the drone’s real-time kinematic positioning using the Virtual Reference Station network from Can-Net (Trimble, Westminster, CO, USA). The GCP positions were measured with a Reach RS2 RTK GNSS receiver (Emlid, Budapest, Hungary), also using the Can-Net network. Overall accuracy was always below 5 cm between dates for the same mission, with an average offset around 2-3 cm. Weather conditions ranged from clear and sunny to cloudy skies with stronger winds for a date (September 14) and a little mist for another (September 22). Weather conditions were noted for each date to analyze the impact of light and wind conditions on the model (Supplementary Table S2).

Once the photos were collected with the drone, they were used to create orthomosaics using Metashape Professional v.1.7.5 (Agisoft, St. Petersburg, Russia). The first step was to calibrate the camera to obtain the best possible accuracy without using GCPs for all the missions (Agisoft, 2021). This was accomplished by doing a drone survey above a field of 0.2 ha with 20 precisely measured GCPs on it. The images were imported into the software and aligned with high precision and generic preselection. GCPs positions were then imported too and moved to align perfectly with the images. The option “Optimize cameras” was used to estimate the best value for the general camera calibration parameters [f, k1, k2, k3, cx, cy, p1 and p2] and the accuracy went from 5 cm to 2 cm. The adjusted values found for the parameters were saved to be used when processing the orthomosaics. This was done by following a protocol largely inspired by Over et al. (2021). Individual photos were imported and aligned with each other using tie points. A major difference is the construction of the digital elevation model (DEM) from the sparse cloud instead of the dense cloud. This choice is motivated by the low flight height, high ground resolution and the species mapped, which are mainly tall grasses with few small shrubs. Under these conditions and using the dense cloud, the software had difficulty aligning individual photographs with the three-dimensional structure of the terrain, and artifacts were created to fill the conflicting information. This problem was greatly reduced when using the sparse point cloud and it allowed us to save time during processing. The orthomosaics were then generated using the previously created DEM. A total of 174 orthomosaics were generated, one for each combination of missions (29) and dates (6).

### 2.4 Annotations dataset

A custom workflow was created to facilitate the annotations of *Phragmites* in the imagery. Annotations were made in the office using the data collected in the field and ArcGIS Pro v.3.0 software (ESRI Inc., Redlands, CA, USA). A grid made up of 128 by 128 pixels squares was created and overlaid on the areas surveyed by the drone. This was done so that the orthomosaics could be cut into tiles of ideal size for training a convolutional neural network, usually a power of two such as 128, 256 or 512. A point, indicating a positive annotation, was added in a distinct map layer inside the grid square where common reed was present. Ground truth data was collected in September to help confirm the species’ presence. Next, using Python and ArcPy, the ArcGIS library for GIS analysis in Python, the orthomosaics were divided into individual raster files representing the grid squares. These extracted tiles were sorted into different folders according to their annotation, positive or negative. This approach made it possible to quickly test the impact of a change in tile size without having to redo all the annotations. For instance, to switch from the original tile size of 128 x 128 px to a doubled size of 256 x 256 px, four tiles were merged before the extraction step. If at least one of the merged tiles had a positive annotation, the resulting tile would also have a positive annotation while the resulting tile would stay a negative annotation if all the merged tiles had negative annotations. The near-perfect alignment of the imagery between the dates also allowed for mutual validation during annotations. For example, the presence of common reed at a specific location in July implied that the species should still be present there in October. The opposite approach could also be applied: the inflorescence of common reed, not visible in early summer and generally appearing between August 3 and August 24, helped us confirm the annotations for July and differentiate *Phragmites* from species with similar vegetative traits, such as reed canary grass. After three dates had been annotated, an active learning approach was applied by training a primitive model to obtain predictions for the remaining dates. It was still necessary to go back over the model predictions at this step to avoid bias in the dataset. Points were added to tiles with Phragmites undetected by the model and removed when corresponding to false positives. Ultimately, there was a total of 45 628 positive annotations, with a progression over the course of the season, from 6635 in July to 7700 on the last date in October.

To test the impact of the change in spatial resolution, imagery was resampled at different GSD values; 0.30, 0.45, 0.60, 0.75, 0.90, 1.20 and 1.50 cm. The last value corresponds to the resolution obtained when flying at 120 m, the maximum altitude permitted by Canadian regulations. Bicubic resampling method was used without the addition of optical blur, giving potentially superior results compared to real-world applications (Brown et al., 2022; Clabaut et al., 2021). The size of the tiles was then resized back to 256 x 256 pixels for all the GSD values to prevent the size of the input from impacting the model results (Richter et al., 2021).

Finaly, before training models, the data was split into a training and a validation dataset. The split was done randomly based on a ratio of 80% and 20% respectively and remained similar throughout the model testing process. Three areas, accounting for approximately 10% of the initial data set, were also kept as the independent test dataset to evaluate model performance on previously unseen data. Overall, there were 176 314, 44 078 and 29 694 tiles in the training, validation, and testing dataset respectively. The complete dataset used will be available soon at https://doi.org/10.20383/103.0954.

### 2.5 Training a model

A CNN was developed to train the model. The code was run using Jupyter Notebook and Python. The model was trained with PyTorch library and its high-level equivalent, fastai. The latter is a library developed by Howard & Gugger (2020a) to simplifies the use of deep learning without in-depth knowledge of computer science (Howard & Gugger, 2020b). All analyses were carried out on a local computer using CUDA and a RTX 4090 graphics processing unit (Nvidia, Santa Clara, CA, USA). Several hyper-parameters were tested to determine the best model. 256 pixels for the tile size, 72 tiles for the batch size, the use of data augmentation, training for 20 epochs, VGG19_BN pre-trained with ImageNet as the architecture, focal loss as loss function and Adam as optimizer gave the best result overall, and all the results presented use these values (Supplementary Table S1).

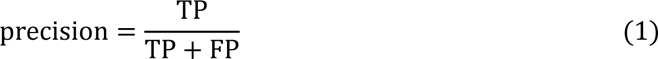

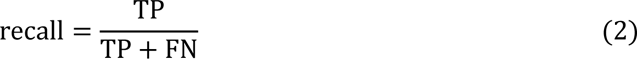

To evaluate the performance of the models, the F1-score was used. This metric is based on the harmonic mean of precision and recall (Equation 3). Precision measures the model’s ability to correctly classify as positive only the examples that are positive, while recall measures the model’s ability to identify all true positive examples. For models with similar F1-Score, recall was favored over precision, as each false negative represent a location where the species is present, and which could become problematic to manage in the longer term if not detected. Therefore, false negatives should be avoided as far as possible. Operational recall and precision metrics were also calculated considering adjacent tiles. Using this calculation method, a false negative became a true positive if at least one of the eight adjacent tiles was a positive annotation, while a false positive remained the same only if none of the adjacent tiles were positive annotations. All results shown in the next section refer to the model inference on the three test zones only: ‘Dense established’, ‘Establishing’ and ‘Post-treatment’.

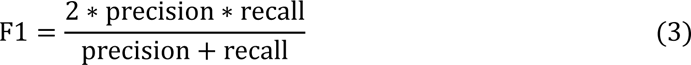

## 3. Results

### 3.1 Over the growing season

The model applied on imagery from either September 14 or 22 achieved the highest F1-scores with value of 0.98, 0.94 and 0.20 for ‘Dense established’, ‘Establishing’ and ‘Post-treatment’ respectively (Table 1). Predictions improved for all three zones over the growing season, whereas the F1-scores for the first date in July were 0.94, 0.75 and 0.08. The lowest recall value was obtained on October 3 for the ‘Dense established’ and ‘Establishing’ zones when light conditions were sunny. The lowest precision value was obtained on July 7 for all three zones while the common reed was still in the vegetative stage. In general, for the ‘Dense established’ zone, metrics were high for all dates, with an average of 0.97 for recall, 0.95 for precision and 0.96 for F1-score. Results were also good for the ‘Establishing’ zone, especially for September imagery, with averages for all dates of 0.84, 0.87 and 0.85 for recall, precision, and F1-score. For the ‘Post-treatment’ zone, recall was generally good, with an average of 0.85, while average precision was only 0.08, reducing the average F1-score at 0.15.

**Table 1.** Evolution of metrics (recall, precision, and F1-score) and operational metrics (recall and precision) for each zone at each date. Best results from the model were obtained on Sept. 22 for the ‘Dense established’ zone and on Sept. 14 for the ‘Establishing’ and ‘Post-treatment’ zone. Also, a significant improvement in results was observed with the operational calculation method, with near-perfect recall for all three zones. This calculation method considers the predictions at the individual level rather than the tile level. The highest recall, precision and F1-score for each zone is shown in bold.

| Zone | Date | Recall | Precision | F1-score | Operational recall | Operational precision |
| --- | --- | --- | --- | --- | --- | --- |
| **Dense established** | **July 07** | 0.99 | 0.90 | 0.94 | 1.00 | 0.98 |
|  | **Aug. 03** | 0.96 | 0.94 | 0.95 | 1.00 | 0.99 |
|  | **Aug. 24** | 0.97 | 0.96 | 0.97 | 1.00 | 1.00 |
|  | **Sept. 14** | 0.96 | 0.96 | 0.96 | 1.00 | 0.99 |
|  | **Sept. 22** | **0.97** | **0.98** | **0.98** | 0.99 | 0.99 |
|  | **Oct. 03** | 0.94 | 0.98 | 0.96 | 1.00 | 0.99 |
| **Establishing** | **July 07** | 0.78 | 0.72 | 0.75 | 1.00 | 0.74 |
|  | **Aug. 03** | 0.85 | 0.78 | 0.81 | 1.00 | 0.78 |
|  | **Aug. 24** | 0.83 | 0.90 | 0.87 | 1.00 | 0.92 |
|  | **Sept. 14** | 0.89 | **0.99** | **0.94** | 1.00 | 1.00 |
|  | **Sept. 22** | **0.92** | 0.91 | 0.92 | 1.00 | 0.95 |
|  | **Oct. 03** | 0.77 | 0.91 | 0.83 | 1.00 | 0.98 |
| **Post-treatment** | **July 07** | 0.84 | 0.04 | 0.08 | 1.00 | 0.09 |
|  | **Aug. 03** | 0.79 | 0.06 | 0.10 | 1.00 | 0.16 |
|  | **Aug. 24** | 0.78 | 0.10 | 0.18 | 1.00 | 0.21 |
|  | **Sept. 14** | 0.92 | **0.11** | **0.20** | 1.00 | 0.25 |
|  | **Sept. 22** | **0.95** | 0.09 | 0.17 | 1.00 | 0.22 |
|  | **Oct. 03** | 0.83 | 0.10 | 0.18 | 0.94 | 0.20 |

The operational calculation method produced near-perfect recall results in all three zones (Table 1), with the total number of combined false negatives decreasing from 170 to just three (one on September 22 in the ‘Dense established’ zone and two on October 3 in the ‘Post-treatment’ zone). The number of false positives was also divided by four in the ‘Dense established’ zone, dropping from 114 to 26. There was a smaller reduction of false positives in the ‘Establishing’ and ‘Post-treatment’ zones, from 44 to 35 and 1852 to 1636 respectively.

We compared annotations and predictions for the three zones using imagery with the best F1-score on heatmaps (Fig. 3). The ‘Dense established’ and ‘Establishing’ zones produced almost identical maps between annotations and model predictions. The area containing common reed is well defined on the prediction maps in both cases. In the case of the ‘Post-treatment’ zone, several false positives are present on the prediction map, making it difficult to accurately target areas where common reed is actually present. Almost all the false positives represent locations where either a species similar to common reed is present, in this case *Phalaris*, or where features that were under-represented in the training dataset, such as tree trunks and branches, are present.

**Figure 1.**
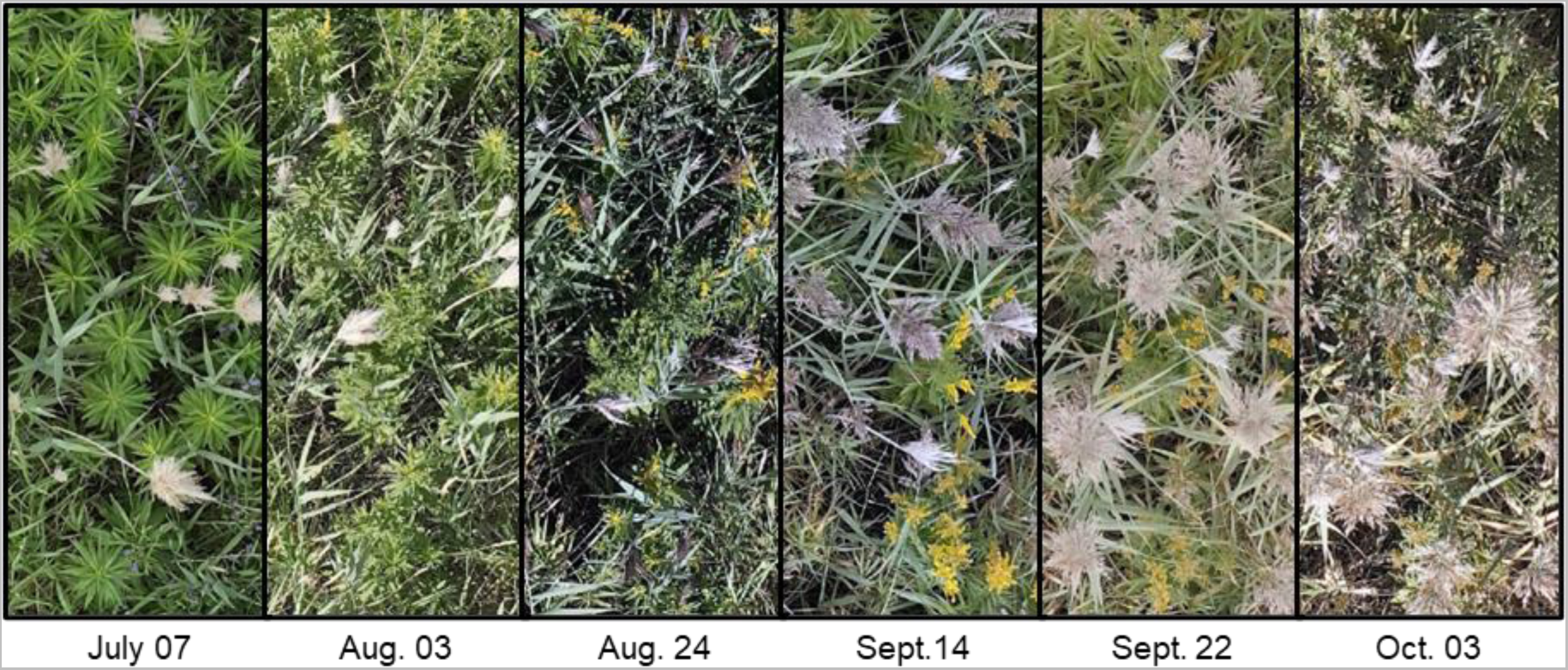
Temporal variations of the same emplacement in the imagery for the 2022 growing season. For this plot, the sky conditions were cloudy on July 7, September 14, and September 22. The sky was sunny for the other dates. Also, the wind was always low (three or less on Beaufort scale), except for September 14 (where it reached five on Beaufort scale).

**Figure 2.**
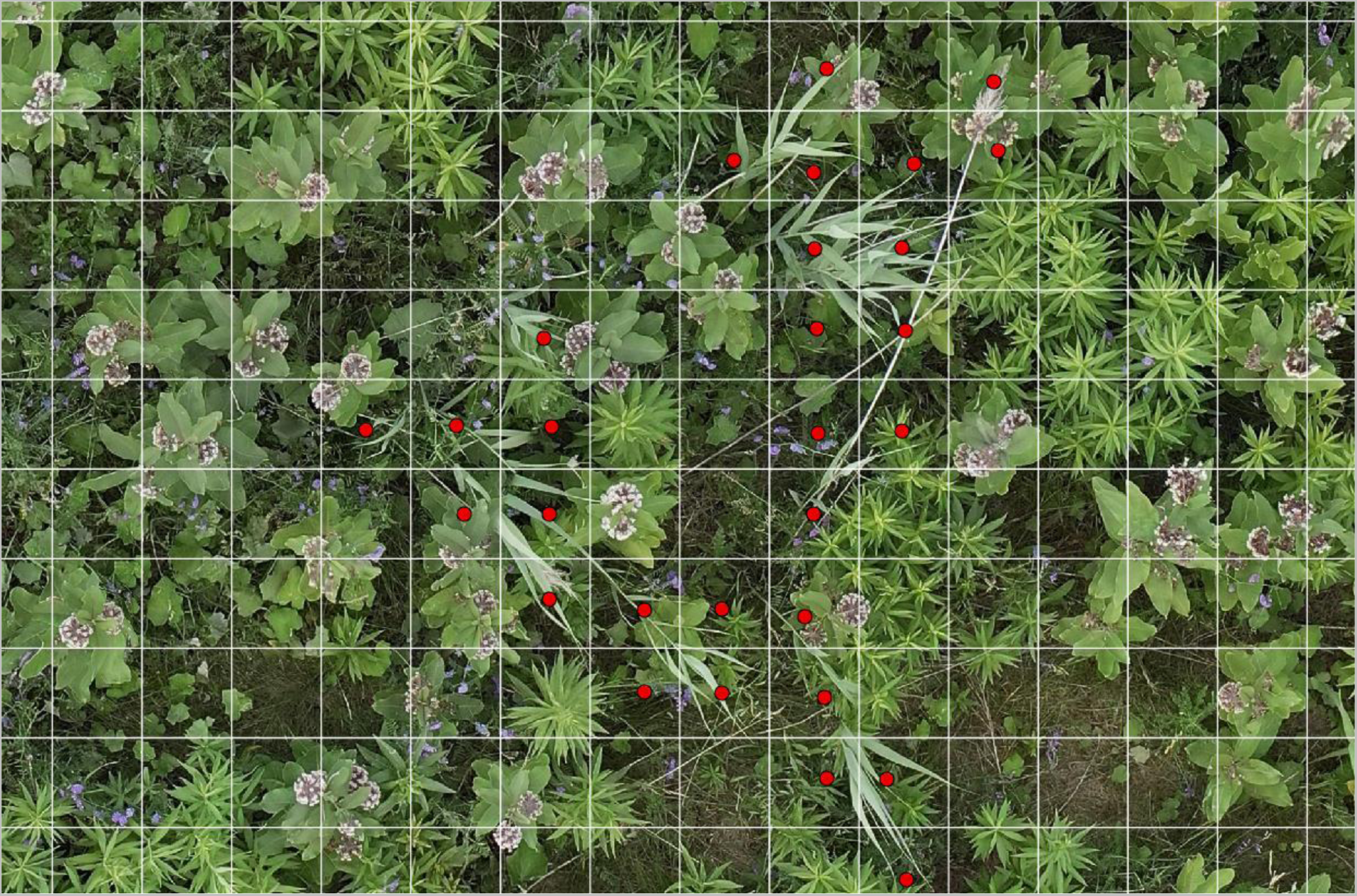
Annotations workflow example with the imagery in background overlaid by a grid and annotation points layer. Each square of the grid has a size of 128 by 128 pixels, representing 19.2 x 19.2 cm at this resolution. The positive annotations where common reed is present are displayed as red dots. In certain instances, discerning the necessity of a positive annotation may pose challenges, particularly when only a leaf tip or inflorescence stem is observed within the square.

**Figure 3.**
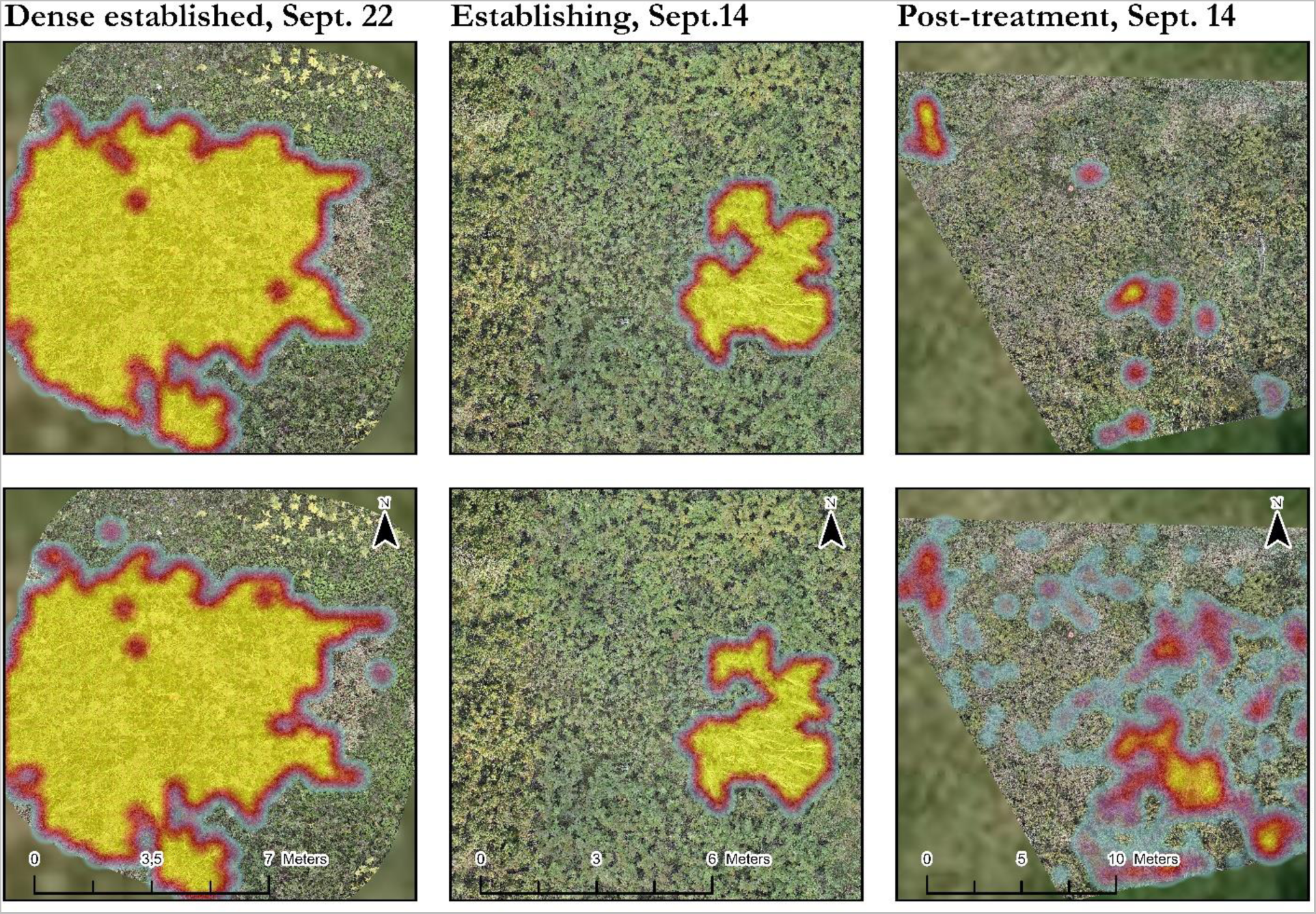
Heat maps of the positive annotations (top) and positive predictions from the model (bottom) for each zone. The maps only show the results for the optimal detection date for each zone. While the annotation and prediction maps are similar for the ‘Dense established’ and ‘Establishing’ zones, ‘Post-treatment’ zone shows a strong presence of almost randomly distributed false positives.

For completeness, we tried to train a model that detects only the seed head, as the inflorescence is a unique feature that seems to favor its detection and could reduce the number of false positives in the post-processing area (Fig. 4). Most stems showed inflorescence from late summer onwards, resulting in similar recall between the two models from this point in the growing season. In terms of model precision, there was a large reduction in false positives with the inflorescence only model, from a total of 1852 for all dates combined to just 68.

**Figure 4.**
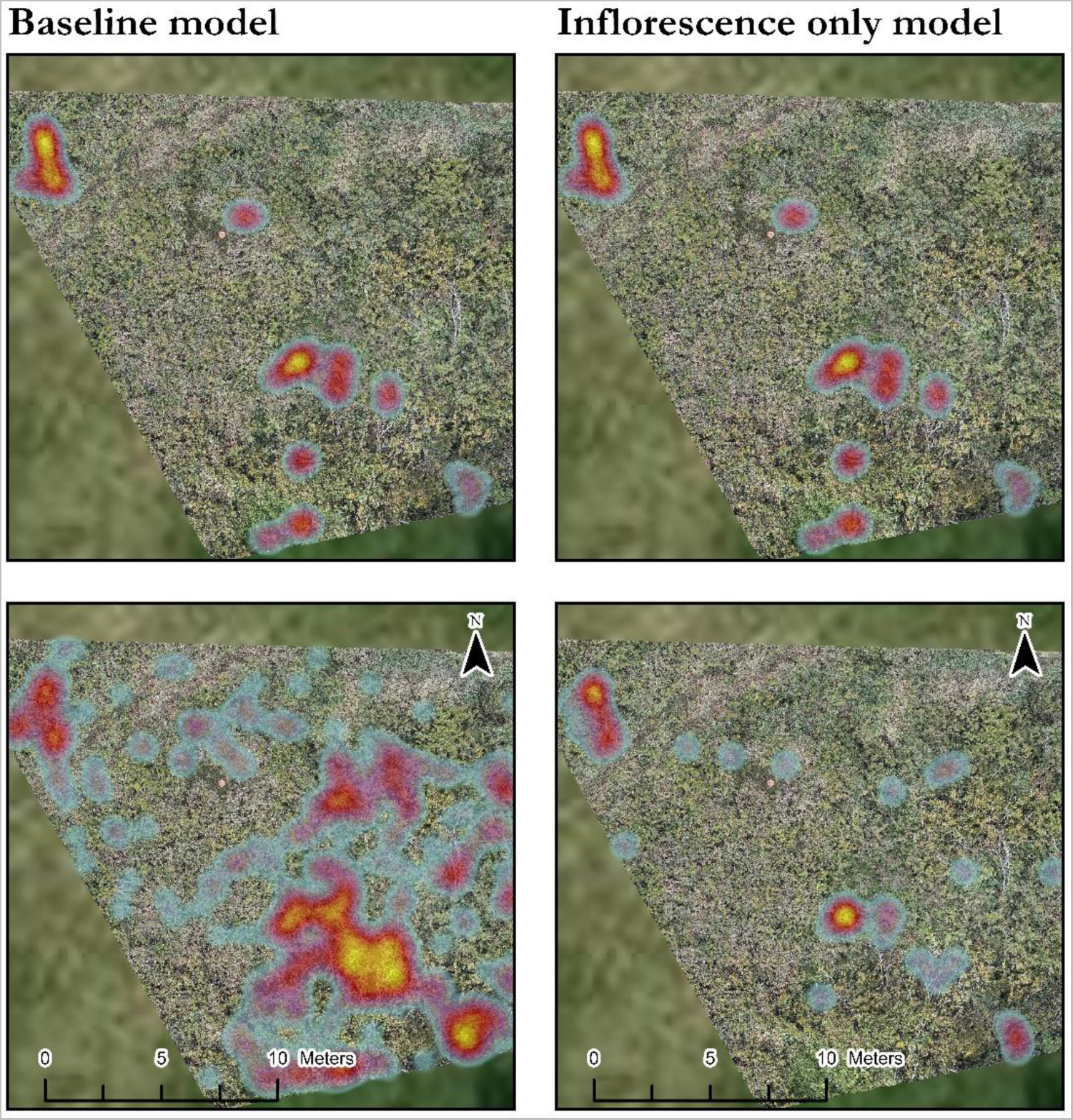
Comparing the baseline model with a model detecting only infloresence, heat maps of the positive annotations (top) and positive predictions from each model (bottom) for the ‘Post-treatment’ zone. The maps only show the results for the optimal detection date for both model (14 Sept.). The baseline predictions are the same as in Fig 3. A model that detects only inflorescence produces a significantly lower number of false positives while still correctly identifying the locations where common reed is present.

The model predictions were better when the common reed to be identified was part of a colony, compared with isolated stems (Fig. 5). Indeed, a higher value of adjacent tiles containing common reed resulted in a better prediction probability. When the tile given to the model was surrounded by tiles also containing *Phragmites*, the probability of a good prediction was 98% (p < 0.0001). As the number of adjacent tiles containing common reed decreased, for instance when reaching the edge of the colony, the probability of a good prediction decreased. In the case of a tile containing an isolated stem of common reed with no adjacent tile containing the species, the probability of a good prediction fell to 63%.

**Figure 5.**
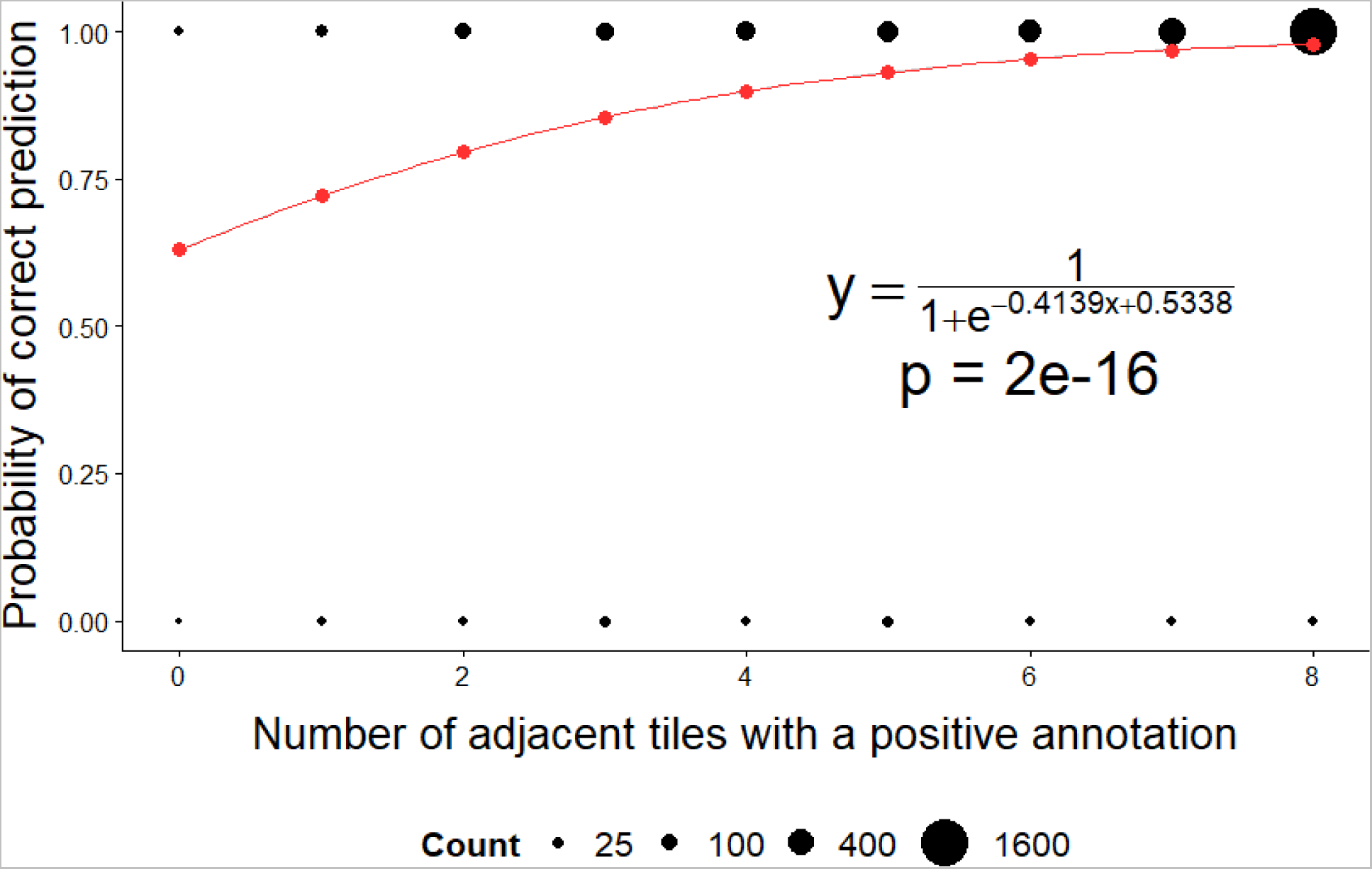
Increase in the probability of a good prediction by the model according to the number of adjacent tiles containing the target species. The count of wrong predictions remained consistent for different number of adjacent tiles containing the target species, ranging from 4 to 27 with a mean of 19 and a median of 21. On the other hand, the count of correct predictions increased exponentially with the number of adjacent tiles containing the target species, rising from 58 to 1634 with a mean of 334 and a median of 153.

### 3.2 Effect of spatial resolution

Testing at different resolutions was carried out on data from September 14 and 22 only for the three zones, as these dates gave the best results over the growing season. For all three zones, there was a trade-off between recall and precision with decreasing spatial resolution (Fig. 6). From an initial resolution of 0.15 cm px^−1^ down to 0.90 cm px^−1^, an increase in precision comes with a decrease in recall, and the opposite is also true depending on the zone. For resolutions of 1.20 and 1.50 cm px^−1^, recall and precision changed in the same direction, but differently for the three zones (constant for the ‘Dense established’ zone, increasing for ‘Establishing’ and decreasing for the ‘Post-treatment’ zone). We also found that the model’s confidence in predictions was generally higher in the case of true positives and true negatives, with mean confidence levels of 90% and 85% respectively for all occurrences (Fig. 7). In contrast, the average model confidence for false negatives was 68%, and 66% for false positives occurrences. The model was therefore more certain of the results when the predictions are correct, and more uncertain in the case of wrong predictions. Changes in model predictions because of changing spatial resolution were not constant and linear. For example, example (b) shows that the model gives a false negative at a resolution of 0.60 and 0.90 cm px^−1^ for this tile, but a lower or higher resolution gives a correct prediction. In the same way, examples (d) and (e) show that false negatives are not always obtained only at low or at high spatial resolution, but rather at any resolution (at 0.15, 0.90 and 1.50 cm px^−1^ in this case).

**Figure 6.**
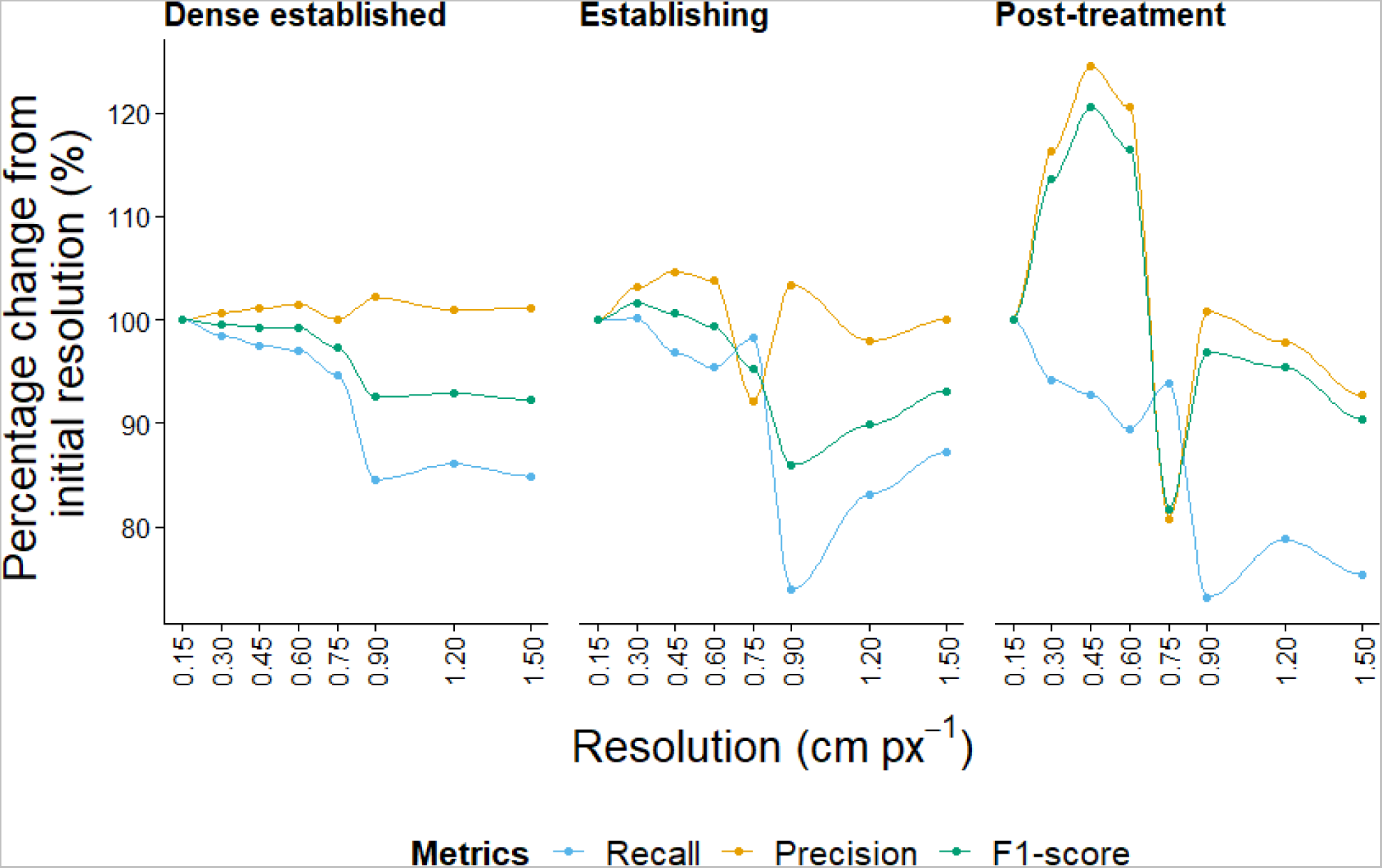
Changes in recall, precision and F1-Score for each zone at different resolutions. A decrease in recall from initial resolution up to 0.90 cm px^−1^ was observed for the three zones, followed by a plateau or slight improvement between a resolution of 0.90 and 1.50 cm px^−1^. Precision showed the opposite result, with an increase from initial resolution up to 0.60 cm px^−1^, then a variable decrease or a plateau between 0.75 and 1.50 cm px^−1^.

**Figure 7.**
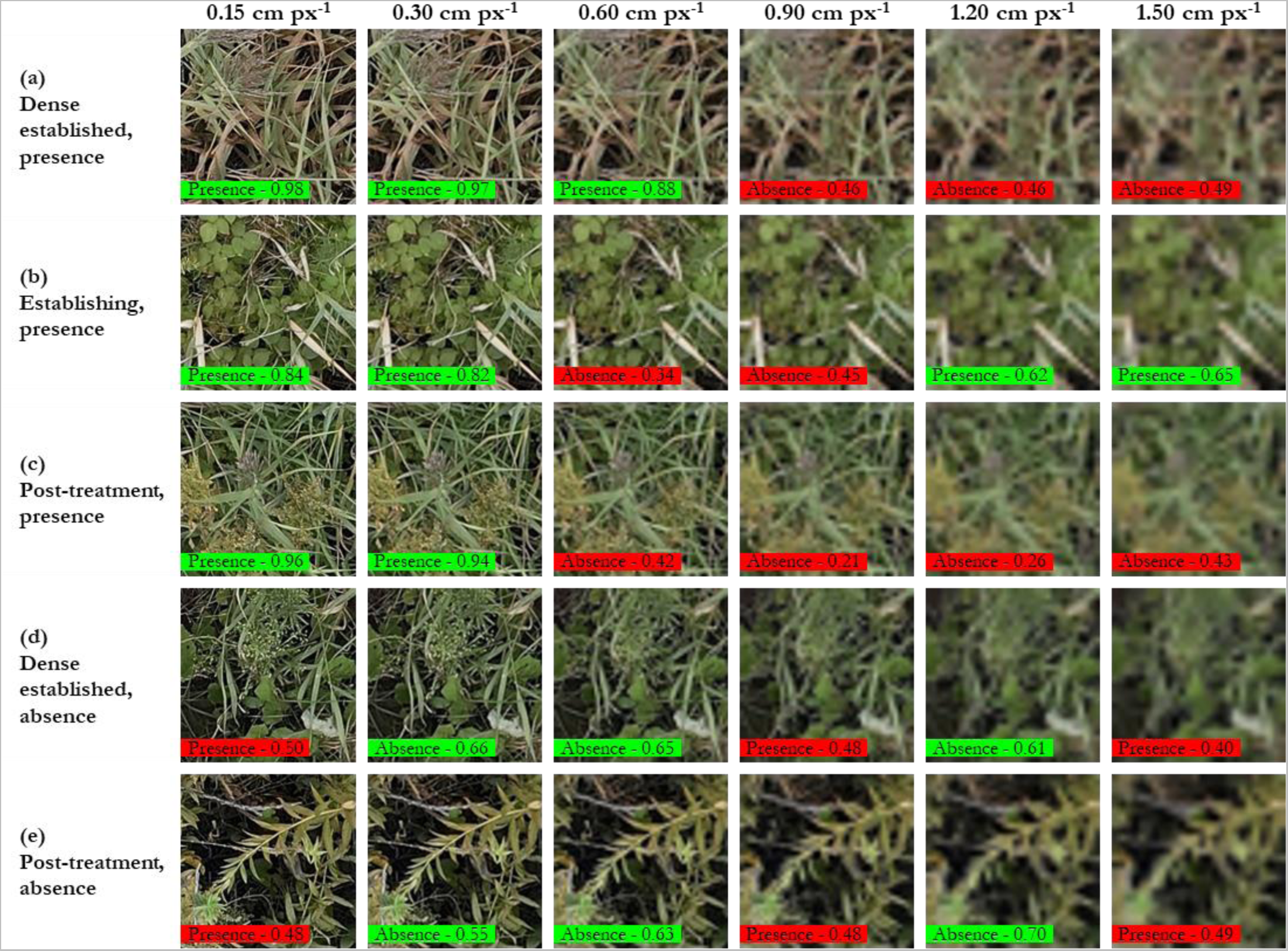
Predictions of the model for various tiles at different resolutions. First three rows show examples of false negatives occurring with a change of spatial resolution, while the last two rows show examples of false positives in the same situation. Color of the prediction box indicates whether the prediction was correct (green) or wrong (red). Confidence of the model is also reported for each tile, where 1.00 and 0 represent perfect confidence, and 0.50 a complete uncertainty of the output. Not all the spatial resolutions tested are shown in the figure.

## 4. Discussion

In this study, we wanted to determine whether early detection of an invasive alien plant was possible with an approach using drone imagery and artificial intelligence. As predicted, our results showed that this strategy was generally successful in detecting common reed within a conservation area. In areas showing a higher degree of invasiveness, our approach detected common reed very well, which is consistent with the literature (Anderson et al., 2021; Cohen & Lewis, 2020; Higgisson et al., 2021). Nevertheless, early detection of common reed in areas where it was actively controlled (i.e. the ‘Post-treatment’ zone) and densities were lower, detection was more difficult, with many false positives. Our results also pointed toward an optimal time for detection, in September, when the inflorescence was clearly visible on the imagery. This supports our hypothesis that unique characteristics enable better identification. We also found that a higher flight altitude up to 60 m (leading to a lower spatial resolution of 0.75 cm pixel^−1^) did not significantly reduce our models’ ability to detect common reed, thereby allowing greater monitoring efficiency by allowing a larger area to be covered in a single flight. Finally, our approach was excellent (in many cases, perfect), when considered from an operational perspective (e.g. when detecting common reed in larger ~1 m^2^ areas). Overall, our study shows high potential for the early detection of common reed using high-resolution RGB drone imagery, but also shows that more work is needed to improve the performance of our model in areas undergoing frequent management to control this species.

### 4.1 Phenology

We found that our ability to detect common reed from drone imagery depended on the time of year. Indeed, our model showed optimal performance when the inflorescence was well-developed and prominently visible in September. We found similar results obtained from the two September dates. This suggests that a model trained solely on these dates would perform well when applied to unseen data taken during this period. In fact, this approach may yield better results than our current method of training a model with all acquisition dates. Models based on two or three dates and applied to this period generally perform better than models trained with just one or, as in our case, models trained on a greater number of dates (Grybas & Congalton, 2021; Weisberg et al., 2021).

Many studies investigating the impact of phenology on remote sensing of plant species with UAVs have focused on forest environments. Nevertheless, these studies have come to the same conclusion that the period when inter-specific differences are most pronounced while intra-specific variation is minimal is the best time to make the identification (Grybas & Congalton, 2021; Hill et al., 2010; Weil et al., 2017). In the case of forest environments, this generally takes place at the start of senescence, during the color change that occurs in autumn in temperate environments (Cloutier et al., 2023). Weisberg et al. (2021) came to a similar conclusion by studying two invasive grasses in a cold desert climate, cheatgrass and medusahead. Ecosystem conditions being different, the best period for detection was May-June and June-July respectively for each species. In both cases, however, this corresponded to the peak seed head production just before the onset of senescence. This shows that flowering can be an important life stage for detecting plant species using drone imagery and computer vision.

In our study, the seed heads of common reed were already becoming visible during the August 24th imagery, which might have helped to achieve similar results as in September. Likewise, the initiation of senescence observed during the October imaging session could have led to results as similar as or even better than those of previous dates. Although the influence of weather conditions on model predictions could not be evaluated separately, it likely influenced our results. Both dates were characterized by sunny skies. Previous studies suggest that variable light conditions, including both cloudy and clear-sky days, can exert some degree of influence on results, although usually to a negligible extent (Dandois et al., 2015; De Souza et al., 2021). Other researchers found that CNNs typically maintain robustness across varied illumination conditions, particularly when employing data augmentation techniques to artificially diversify the brightness conditions of the training dataset (Barbedo et al., 2019; Lisein et al., 2015; Safonova et al., 2019).

Elevated exposure levels can still pose challenges in distinguishing between various color tones and the presence of large dark shadows can result in an elevated occurrence of omission errors (Lopatin et al., 2019; Milas et al., 2017). This problem was observed even prior to model training when annotating the imagery, with sunny days causing more difficulties. Additionally, it may also prevent the identification of smaller individuals, which find themselves surrounded by taller species that overshadow them. This could potentially account for the observed variation in recall between different dates, particularly in the ‘Establishing’ and ‘Post-treatment’ zones compared to the ‘Dense Established’ zone. In the latter, shadows are primarily cast by neighboring common reed individuals within the monospecific colony, facilitating accurate model predictions. Conversely, in the other two zones, the presence of adjacent different species can obscure less dominant common reed specimens with their shadows. Increasing the frequency of data collection could help to mitigate the impact of diverse weather conditions. However, despite training the model under highly variable conditions, optimal results were obtained in ideal conditions, such as cloudy weather with diffused light. Also, while moderate to low wind conditions may enhance drone operability, their influence on data acquisition was minimal. The imageries were not blurrier on windy days (Sept. 14 and 22) and the impact of wind on results seems negligible, which is in line with existing literature (Dandois et al., 2015).

Moreover, in the case of common reed, as with other invasive alien species, the species generally has a longer growing season (Vymazal & Krőpfelová, 2005; Wolkovich & Cleland, 2011). Consequently, one might have anticipated enhanced detection performance in October. However, analysis of imagery from October, coinciding with the onset of senescence marked by the progressive yellowing of leaves in various species yielded suboptimal results compared to September. This discrepancy can be attributed to several factors discussed earlier, particularly regarding the sunny-sky conditions in the October imagery. Additionally, common reed begins its growth earlier in the season than our earliest imagery, and it would have been interesting to start data acquisition at the beginning of the growing season. However, it is uncertain whether the results would have been favorable. Confused species reed canary grass can emerge up to a month earlier than common reed in temperate climates, helping to achieve greater aboveground biomass earlier in the season (Vymazal & Krőpfelová, 2005). It could have led to more confusion by the model without allowing for species differentiation. These species can exhibit comparable phenological characteristics at some period in the growing season, such as invasive early-season grasses, which might reduce model performance, given the higher likelihood of confusion among them (Bradley, 2014).

### 4.2 Between the zones

As expected, we obtained better results in areas with a higher degree of common reed invasion than in areas managed to control the species. Common reed individuals are more readily identifiable by the model when they are surrounded by other reed individuals, within a colony for example. For instance, in the ‘Dense Established’ and ‘Establishing’ zones, the maps for the optimal prediction time, in September, are very similar between annotations and model predictions. By contrast, individuals located at the periphery of the colony and isolated stems with limited or absent nearby common reed were less well detected by the model. Lower results were obtained in actively controlled zone, which is comparable to the use of multispectral or hyperspectral imageries for detection, albeit for distinct reasons. While our approach using UAVs exhibits reduced efficacy, unlike hyperspectral imagery, the challenge does not arise from an incapacity to detect the individual stems of common reed (Elmer et al., 2021; Pengra et al., 2007). When using hyperspectral or multispectral imageries, the low spatial resolution is frequently the cause of mixed pixels, whereby the common reed is in minority within the pixel and its spectral signature is insufficient to allow for its detection (Hsieh et al., 2001). In our study, isolated stems were generally well detected, as evidenced by the high recall of 0.92 for both September dates in the ‘Post-treatment’ zone.

Our approach was limited by the very large number of false positives in the ‘Post-treatment’ site. This would limit its usefulness in practice, given the significant time cost of having to travel to these locations in the field. However, in the case of an invasive alien plant, an overprediction (high recall, low precision) is preferable to avoid the species becoming established if it is not adequately treated. However, if the costs of prevention are higher than the benefits due to many false positives, the approach may not be ideal (Jiménez-Valverde et al., 2011). One of the reasons for the large number of false positives in the ‘Post-treatment’ area could be the discrepancy between the test and training datasets. A cluster of small, semi-dormant trees was present in the middle of the testing zone, with several tree branches clearly visible. These features were not present in the training set, so the model incorrectly predicted it as common reed. It would have been necessary to conduct some fine-tuning to ensure that the model had been exposed to these features during training and could differentiate them from those characteristic of the common reed (Han et al., 2019). Another contributing factor was the greater prevalence of reed canary grass, a species that closely resembles reed in its vegetative stage. Reed canary grass can exhibit invasive characteristics but is not treated by park staff, which explains its high abundance in the area. Given the more pronounced class imbalance observed in this zone, with a low number of true positives for common reed, the model generated a greater number of false positives by incorrectly identifying the reed canary grass as common reed. Also, given the low presence of common reed following the annual treatment, it is more akin to detecting a rare species, which generally yields poorer results due to its intrinsically lower presence (Leduc & Knudby, 2018). One potential solution to this issue could be the use of environmental variables to more effectively identify potential establishment sites and disregard those deemed implausible (Reckling et al., 2021). For instance, the distribution patterns of common reed are relatively well documented, and the use of a digital surface model (DSM) or topographic moisture index (TMI) could enhance the precision of location prediction.

In order to reduce the number of false positives in the ‘Post-treatment’ site, we also tried to train a model that focused only on seedhead detection. The inflorescence is usually visible in reeds from the second to the fourth growing season after germination (Haslam, 1971). Nevertheless, despite treatment against the species in the autumn preceding the field period, a seed head was visible on almost all common reed stems in this area from mid-August onwards. Given that the number of false positives fell by 95% while recall remained stable, this could represent an effective avenue for detecting the species efficiently in management areas without incurring additional costs with the presence of numerous false positives to check. One limitation of this approach is that common reed plants must have an inflorescence to be detected by the model. This would not be the case with young seedlings from seed germination. If the inflorescence only appears the following season and the plant is not detected, this will leave an extra growing season for the plant to establish, which could complicate eradication. Furthermore, an approach that only identifies inflorescences would need to be used from late August onwards, not earlier, to ensure that the inflorescence is well developed at time of imagery acquisition. The time of inflorescence development could also vary from year to year. Therefore, it would be essential to ensure that imagery is acquired after this period.

### 4.3 Spatial resolution

It can be observed that, in general, a reduction in spatial resolution leads to a decline in the number of positive predictions made by the model. This has implications for both precision and recall. An increase in precision is linked to fewer false positives, while a decrease in recall is linked with a higher number of false negatives. In our study, we found that the decline in recall was minimal up to a GSD of 0.75 cm, after which it declined rapidly for all three zones. Precision remained relatively stable across all spatial resolutions, exhibiting minimal variation. The ‘Post-treatment’ zone showed greater variability in results, which could be attributed to the representation of data in percentage of change relative to performance measurements at the initial resolution. Initially, precision in this area was relatively low (0.10), suggesting that a change of a few percentage points may have a more pronounced impact compared to zones like ‘Dense Established’ and ‘Establishing’, where initial values were already high (>0.90). It is conceivable that flying higher would not have significantly affected the outcome. In a study on remote sensing of cattle, a target larger than the common reed, it was observed that between GSD values of 1, 2, and 4 cm, the intermediate value yielded superior results when training different CNNs (Barbedo et al., 2019). This can be attributed to the fact that an excessively high spatial resolution prevents the optimal extraction of crucial characteristics of the target species. Other studies have shown that a spatial resolution of 3 to 5 cm px^−1^ is sufficient to detect most plants and differentiate communities (Kattenborn, Eichel, et al., 2019).

Considering significant limitations that may artificially improve results, such as the lack of optical blur, it is uncertain whether a flight altitude of 60 m, can be definitively deemed ideal (Brown et al., 2022; Clabaut et al., 2021). The absence of optical blur might have led to unrealistically sharp degraded images due to pixel subsampling compared with actual flights at higher altitudes. Thus, it would have been better to repeat the experiment with imagery taken at different altitudes, without using artificial resampling to simulate a reduction in spatial resolution. Although our results are likely to be more optimistic than the reality in the field, we can still assume that the optimum flying height lies between 15 and 60 meters, not higher. Furthermore, the linear trend between spatial resolution and performance measurements should probably be similar in the case of imagery actually acquired at different altitudes. Consequently, we anticipate observing a comparable initial plateau, where a decrease in spatial resolution minimally affects results, followed by a threshold value beyond which recall declines rapidly. This inflection point is anticipated to manifest earlier, potentially around a spatial resolution of 0.30 or 0.45 cm px^−1^, corresponding to flights at 24 and 36 m respectively.

### 4.4 Operationnal metrics

The annotations and predictions were made at the tile level, which may not be the most relevant spatial scale in an operational field setting. A team of biologists surveying the field would not be constrained by these tile sizes, which only represent a quadrat of approximately 40 by 40 cm at the highest resolution. Therefore, we used an alternative method for comparing the different approaches. In our study, the model does not have access to contextual information when making predictions, as it processes one tile at a time, independently from each other. For comparison, when annotating, annotators could see the context (i.e. neighbourhood), which facilitated annotation. The model sometimes misidentifies bits of reed or identifies them as present when they are not, resulting in false positives. Similarly, if a tile containing reed is not detected by the model, it may result in a false negative. Nevertheless, if at least one adjacent tile also contains common reed and is well predicted by the model, from an operational point of view there are no false negatives in this case. Consequently, we considered adjacent tiles to obtain performance measurements at the level of reed individuals or the usual one-meter quadrat, rather than at the tile level alone. A considerable increase in recall was observed in this context, indicating that the predictions are highly accurate in terms of recall. This enables the species to be eradicated before it can become established.

There may also be variability between different biologists who must identify the species in the field, which may also result in some instances of omission in the current field survey method (Morrison, 2016). For the ‘Dense Established’ and ‘Establishing’ zones, an increase in precision was also observed to achieve near-perfect values in September, the best time for detection. This suggests that many false positives were possibly annotation errors with bits of common reed in the tile that were not determined to be sufficiently visible to be considered as common reed presence by the annotators. The only area where precision remained low despite the calculation including adjacent tiles was in the ‘Post-treatment’ zone. In that zone, the high presence of false positives was therefore not linked to the actual presence of common reed in the vicinity, but rather to other factors such as the presence of reed canary grass and tree branches, as discussed above. The use of a model detecting only the inflorescence could therefore be preferable in this area, despite the possible limitations.

In conclusion, our study confirms that an approach using drones and artificial intelligence is effective in detecting common reed, especially in autumn. We only tested one invasive alien plant in a specific location as part of this project, and it would be worthwhile to apply this technique to other invasive alien plant species and in diverse environments. Also, generating the dataset poses a significant challenge due to the time-intensive nature of its construction, compounded by the necessity for a sufficiently extensive dataset to achieve precise outcomes (Kattenborn et al., 2021). Hence, the data, models, and scripts developed in this project, encompassing imagery and annotations of common reed presence, are openly accessible, aiming to foster collaboration and alleviate the burden of dataset creation. Furthermore, the aim was not necessarily to achieve perfection in results, as technology is advancing rapidly, and new approaches are already on the rise such as vision transformers. Such techniques, including those demanding reduced supervision during training, could hold even higher promise for yielding improved near-term results in remote sensing of invasive alien plant species.

## Acknowledgments

The authors thank Sabrina Demers-Thibault, Charles Picard-Krashevski, Léo Benoit-Charest and Guillaume Tougas for their contribution to the fieldwork for this study. Furthermore, the authors thank the Société des établissements de plein air du Québec (Sépaq), especially Sophie Tessier and Nathalie Rivard from the Parc national des Îles-de-Boucherville, for granting us access and sampling authorization at our study site, and Sam Karathanos from Cambium Phytotechnologies for giving us GIS data. Finally, the authors thank all LEFO lab member and IRBV colleagues for their enriching discussions and advice.

## Funding

A.C.G. received scholarships from the Natural Sciences and Engineering Research Council of Canada (NSERC), from the Fond de recherche du Québec - Nature et Technologies (FRQNT), from Hydro-Québec, and from the Université de Montréal. This study was also funded by the Canada First Research Excellence Fund to the Institute for Data Valorisation (IVADO, Grant number PRF-2021-02), a Discovery grant from the Natural Science and Engineering Research Council of Canada to E.L. (NSERC, Grant number RGPIN-2019-04537) and from the Canada Research Chair in Plant Functional Biodiversity to E.L. (Grant number 950-231858).

## Notes

https://doi.org/10.20383/103.0954

### Competing Interest Statement

The authors have declared no competing interest.

## Références bibliographiques

Agisoft. (2021). *Agisoft Metashape User Manual—Professional Edition, Version 1.7*. https://www.agisoft.com/pdf/metashape-pro_1_7_en.pdf

Aleissaee, A. A., Kumar, A., Anwer, R. M., Khan, S., Cholakkal, H., Xia, G.-S., & Khan, F. S. (2023). Transformers in Remote Sensing: A Survey. Remote Sensing, 15(7), Article 7. 10.3390/rs15071860

Allison, S. D., & Vitousek, P. M. (2004). Rapid nutrient cycling in leaf litter from invasive plants in Hawaii. Oecologia, 141(4), 612–619. 10.1007/s00442-004-1679-z

Alpert, P., Bone, E., & Holzapfel, C. (2000). Invasiveness, invasibility and the role of environmental stress in the spread of non-native plants. *Perspectives in Plant Ecology*, Evolution and Systematics, 3(1), 52–66. 10.1078/1433-8319-00004

Anderson, C. J., Heins, D., Pelletier, K. C., Bohnen, J. L., & Knight, J. F. (2021). Mapping Invasive Phragmites australis Using Unoccupied Aircraft System Imagery, Canopy Height Models, and Synthetic Aperture Radar. Remote Sensing, 13(16), 3303. 10.3390/rs13163303

Baillie, J., Hilton-Taylor, C., & Stuart, S. N. (Eds.). (2004). 2004 IUCN red list of threatened species: A global species assessment. IUCN--The World Conservation Union.

Barbedo, J. G. A., Koenigkan, L. V., Santos, T. T., & Santos, P. M. (2019). A Study on the Detection of Cattle in UAV Images Using Deep Learning. Sensors, 19(24), 5436. 10.3390/s19245436

Barnosky, A. D., Matzke, N., Tomiya, S., Wogan, G. O. U., Swartz, B., Quental, T. B., Marshall, C., McGuire, J. L., Lindsey, E. L., Maguire, K. C., Mersey, B., & Ferrer, E. A. (2011). Has the Earth’s sixth mass extinction already arrived? Nature, 471(7336), 51–57. 10.1038/nature09678

Bazi, Y., Bashmal, L., Rahhal, M. M. A., Dayil, R. A., & Ajlan, N. A. (2021). Vision Transformers for Remote Sensing Image Classification. Remote Sensing, 13(3), Article 3. 10.3390/rs13030516

Bellard, C., Cassey, P., & Blackburn, T. M. (2016). Alien species as a driver of recent extinctions. Biology Letters, 12(2), 20150623. 10.1098/rsbl.2015.0623

Beugnon, R., Le Guyader, N., Milcu, A., Lenoir, J., Puissant, J., Morin, X., & Hättenschwiler, S. (2024). Microclimate modulation: An overlooked mechanism influencing the impact of plant diversity on ecosystem functioning. Global Change Biology, 30(3), e17214. 10.1111/gcb.17214

Bradley, B. A. (2014). Remote detection of invasive plants: A review of spectral, textural and phenological approaches. Biological Invasions, 16(7), 1411–1425. 10.1007/s10530-013-0578-9

Brisson, J., Paradis, É., & Bellavance, M.-È. (2008). Evidence of Sexual Reproduction in the Invasive Common Reed (Phragmites australis subsp. australis; Poaceae) in Eastern Canada: A Possible Consequence of Global Warming. Rhodora, 110(942), 225–230. 10.3119/07-15.1

Brodrick, P. G., Davies, A. B., & Asner, G. P. (2019). Uncovering Ecological Patterns with Convolutional Neural Networks. Trends in Ecology & Evolution, 34(8), 734–745. 10.1016/j.tree.2019.03.006

Brooks, W. R., Lockwood, J. L., & Jordan, R. C. (2013). Tropical paradox: A multi-scale analysis of the invasion paradox within Miami Rock Ridge tropical hardwood hammocks. Biological Invasions, 15(4), 921–930. 10.1007/s10530-012-0340-8

Brown, J., Qiao, Y., Clark, C., Lomax, S., Rafique, K., & Sukkarieh, S. (2022). Automated aerial animal detection when spatial resolution conditions are varied. Computers and Electronics in Agriculture, 193, 106689. 10.1016/j.compag.2022.106689

Byun, C., de Blois, S., & Brisson, J. (2013). Plant functional group identity and diversity determine biotic resistance to invasion by an exotic grass. Journal of Ecology, 101(1), 128–139. 10.1111/1365-2745.12016

Byun, C., de Blois, S., & Brisson, J. (2018). Management of invasive plants through ecological resistance. Biological Invasions, 20(1), 13–27. 10.1007/s10530-017-1529-7

Canadian Food Inspection Agency. (2008). *Invasive Alien Plants in Canada—Technical Report*. https://epe.lac-bac.gc.ca/100/206/301/cfia-acia/2011-09-21/www.inspection.gc.ca/english/plaveg/invenv/techrpt/techrese.shtml

Catling, P. M., & Mitrow, G. (2005). A Prioritized List of the Invasive Alien Plants of Natural Habitats in Canada. Canadian Botanical Association Bulletin, 38, 55–57.

Cavender-Bares, J., Schneider, F. D., Santos, M. J., Armstrong, A., Carnaval, A., Dahlin, K. M., Fatoyinbo, L., Hurtt, G. C., Schimel, D., Townsend, P. A., Ustin, S. L., Wang, Z., & Wilson, A. M. (2022). Integrating remote sensing with ecology and evolution to advance biodiversity conservation. Nature Ecology & Evolution, 6(5), 506–519. 10.1038/s41559-022-01702-5

Ceballos, G., Ehrlich, P. R., Barnosky, A. D., García, A., Pringle, R. M., & Palmer, T. M. (2015). Accelerated modern human–induced species losses: Entering the sixth mass extinction. Science Advances, 1(5), e1400253. 10.1126/sciadv.1400253

Clabaut, É., Lemelin, M., Germain, M., Bouroubi, Y., & St-Pierre, T. (2021). Model Specialization for the Use of ESRGAN on Satellite and Airborne Imagery. Remote Sensing, 13(20), 4044. 10.3390/rs13204044

Clevering, O. A., & Van Der Toorn, J. (2000). Observations on the colonization of a young polder area in the netherlands with special reference to the clonal expansion ofPhragmites australis. Folia Geobotanica, 35(4), 375–387. 10.1007/BF02803550

Cloutier, M., Germain, M., & Laliberté, E. (2023). Influence of Temperate Forest Autumn Leaf Phenology on Segmentation of Tree Species from UAV Imagery Using Deep Learning. 10.1101/2023.08.03.548604

Cohen, J. G., & Lewis, M. J. (2019). Development of an Automated Monitoring Platform for Invasives in Coastal Ecosystems. 10.13140/RG.2.2.26336.43522

Cohen, J. G., & Lewis, M. J. (2020). Development of an Automated Monitoring Platform for Invasive Plants in a Rare Great Lakes Ecosystem Using Uncrewed Aerial Systems and Convolutional Neural Networks*. 2020 International Conference on Unmanned Aircraft Systems (ICUAS), 1553–1564. 10.1109/ICUAS48674.2020.9214035

Colautti, R. I., Bailey, S. A., van Overdijk, C. D. A., Amundsen, K., & MacIsaac, H. J. (2006). Characterised and Projected Costs of Nonindigenous Species in Canada. Biological Invasions, 8(1), 45–59. 10.1007/s10530-005-0236-y

Culliney, T. W. (2005). Benefits of Classical Biological Control for Managing Invasive Plants. Critical Reviews in Plant Sciences, 24(2), 131–150. 10.1080/07352680590961649

Dandois, J., Olano, M., & Ellis, E. (2015). Optimal Altitude, Overlap, and Weather Conditions for Computer Vision UAV Estimates of Forest Structure. Remote Sensing, 7(10), 13895– 13920. 10.3390/rs71013895

Davies, K. F., Chesson, P., Harrison, S., Inouye, B. D., Melbourne, B. A., & Rice, K. J. (2005). Spatial Heterogeneity Explains the Scale Dependence of the Native-Exotic Diversity Relationship. Ecology, 86(6), 1602–1610. 10.1890/04-1196

De Souza, R., Buchhart, C., Heil, K., Plass, J., Padilla, F. M., & Schmidhalter, U. (2021). Effect of Time of Day and Sky Conditions on Different Vegetation Indices Calculated from Active and Passive Sensors and Images Taken from UAV. Remote Sensing, 13(9), 1691. 10.3390/rs13091691

Demertzis, K., & Iliadis, L. (2017). Adaptive Elitist Differential Evolution Extreme Learning Machines on Big Data: Intelligent Recognition of Invasive Species. In P. Angelov, Y. Manolopoulos, L. Iliadis, A. Roy, & M. Vellasco (Eds.), Advances in Big Data (Vol. 529, pp. 333–345). Springer International Publishing. 10.1007/978-3-319-47898-2_34

Deng, J., Dong, W., Socher, R., Li, L.-J., Kai Li, & Li Fei-Fei. (2009). ImageNet: A large-scale hierarchical image database. 2009 IEEE Conference on Computer Vision and Pattern Recognition, 248–255. 10.1109/CVPR.2009.5206848

Ding, R., Luo, J., Wang, C., Yu, L., Yang, J., Wang, M., Zhong, S., & Gu, R. (2023). Identifying and mapping individual medicinal plant Lamiophlomis rotata at high elevations by using unmanned aerial vehicles and deep learning. Plant Methods, 19(1), 38. 10.1186/s13007-023-01015-z

Drake, S. J., Weltzin, J. F., & Parr, Patric. D. (2003). *Assessment of Non-Native Invasive Plant Species on the United States Department of Energy Oak Ridge National Environmental Research Park*.

Ehrenfeld, J. G. (2010). Ecosystem Consequences of Biological Invasions. Annual Review of Ecology, Evolution, and Systematics, 41(1), 59–80. 10.1146/annurev-ecolsys-102209-144650

Elmer, K., Kalacska, M., & Arroyo-Mora, J. P. (2021). Mapping the Extent of Invasive Phragmites australis subsp. Australis From Airborne Hyperspectral Imagery. Frontiers in Environmental Science, 9, 757871. 10.3389/fenvs.2021.757871

Evette, A., Breton, V., Petit, A., Dechaume-Moncharmont, C., & Brasier, W. (2019). Les techniques de bâchage pour le contrôle de la renouée. Revue Science Eaux & Territoires, 6. 10.14758/SET-REVUE.2019.1.11

Fridley, J. D., Brown, R. L., & Bruno, J. F. (2004). Null Models of Exotic Invasion and Scale-Dependent Patterns of Native and Exotic Species Richness. Ecology, 85(12), 3215–3222. 10.1890/03-0676

Fridley, J. D., Stachowicz, J. J., Naeem, S., Sax, D. F., Seabloom, E. W., Smith, M. D., Stohlgren, T. J., Tilman, D., & Holle, B. V. (2007). The invasion paradox: Reconciling pattern and process in species invasions. Ecology, 88(1), 3–17. 10.1890/0012-9658(2007)88[3:TIPRPA]2.0.CO;2

Garcin, C., Joly, A., Bonnet, P., Affouard, A., Jean-Christophe Lombardo, Chouet, M., Servajean, M., Titouan Lorieul, & Salmon, J. (2021). *Pl@ntNet-300K image dataset* (1.1) [dataset]. [object Object]. 10.5281/ZENODO.5645731

Giakoumi, S., Katsanevakis, S., Albano, P. G., Azzurro, E., Cardoso, A. C., Cebrian, E., Deidun, A., Edelist, D., Francour, P., Jimenez, C., Mačić, V., Occhipinti-Ambrogi, A., Rilov, G., & Sghaier, Y. R. (2019). Management priorities for marine invasive species. Science of The Total Environment, 688, 976–982. 10.1016/j.scitotenv.2019.06.282

Gigante, D., Venanzoni, R., & Zuccarello, V. (2011). Reed die-back in southern Europe? A case study from Central Italy. Comptes Rendus Biologies, 334(4), 327–336. 10.1016/j.crvi.2011.02.004

Gilbert, B., & Levine, J. M. (2013). Plant invasions and extinction debts. Proceedings of the National Academy of Sciences of the United States of America, 110(5), 1744–1749. 10.1073/pnas.1212375110

Gormley, A. M., Forsyth, D. M., Griffioen, P., Lindeman, M., Ramsey, D. S. L., Scroggie, M. P., & Woodford, L. (2011). Using presence-only and presence–absence data to estimate the current and potential distributions of established invasive species. Journal of Applied Ecology, 48(1), 25–34. 10.1111/j.1365-2664.2010.01911.x

Grybas, H., & Congalton, R. G. (2021). A Comparison of Multi-Temporal RGB and Multispectral UAS Imagery for Tree Species Classification in Heterogeneous New Hampshire Forests. Remote Sensing, 13(13), 2631. 10.3390/rs13132631

Guo, Q. (2015). No consistent small-scale native–exotic relationships. Plant Ecology, 216(9), 1225–1230. 10.1007/s11258-015-0503-7

Han, T., Liu, C., Yang, W., & Jiang, D. (2019). Learning transferable features in deep convolutional neural networks for diagnosing unseen machine conditions. ISA Transactions, 93, 341–353. 10.1016/j.isatra.2019.03.017

Haslam, S. M. (1971). The Development and Establishment of Young Plants of Phragmites communis Trin. Annals of Botany, 35(143), 1059–1072.

Herbold, B., & Moyle, P. B. (1986). Introduced Species and Vacant Niches. The American Naturalist, 128(5), 751–760. 10.1086/284600

Higgisson, W., Cobb, A., Tschierschke, A., & Dyer, F. (2021). Estimating the cover of *PHRAGMITES AUSTRALIS* using unmanned aerial vehicles and neural networks in a semi-arid wetland. River Research and Applications, rra.3832. 10.1002/rra.3832

Hill, R. A., Wilson, A. K., George, M., & Hinsley, S. A. (2010). Mapping tree species in temperate deciduous woodland using time-series multi-spectral data. Applied Vegetation Science, 13(1), 86–99. 10.1111/j.1654-109X.2009.01053.x

Hoeser, T., & Kuenzer, C. (2020). Object Detection and Image Segmentation with Deep Learning on Earth Observation Data: A Review-Part I: Evolution and Recent Trends. Remote Sensing, 12(10), 1667. 10.3390/rs12101667

Howard, J., & Gugger, S. (2020a). Deep learning for coders with fastai and PyTorch: AI applications without a PhD (First edition). O’Reilly.

Howard, J., & Gugger, S. (2020b). Fastai: A Layered API for Deep Learning. Information, 11(2), Article 2. 10.3390/info11020108

Hsieh, P.-F., Lee, L. C., & Nai-Yu Chen. (2001). Effect of spatial resolution on classification errors of pure and mixed pixels in remote sensing. IEEE Transactions on Geoscience and Remote Sensing, 39(12), 2657–2663. 10.1109/36.975000

Huang, C., & Asner, G. (2009). Applications of Remote Sensing to Alien Invasive Plant Studies. Sensors, 9(6), 4869–4889. 10.3390/s90604869

Hudon, C., Gagnon, P., & Jean, M. (2005). Hydrological factors controlling the spread of common reed (*Phragmites australis*) in theSt. Lawrence River (Québec, Canada). Écoscience, 12(3), 347–357. 10.2980/i1195-6860-12-3-347.1

Hulme, P. E. (2008). Contrasting alien and native plant species-area relationships: The importance of spatial grain and extent. Global Ecology and Biogeography, 17(5), 641– 647. 10.1111/j.1466-8238.2008.00404.x

Huylenbroeck, L., Laslier, M., Dufour, S., Georges, B., Lejeune, P., & Michez, A. (2020). Using remote sensing to characterize riparian vegetation: A review of available tools and perspectives for managers. Journal of Environmental Management, 267, 110652. 10.1016/j.jenvman.2020.110652

IPBES. (2019). *The global assessment report of the intergovernmental science-policy platform on biodiversity and ecosystem services* (E. S. Brondízio, J. Settele, S. Díaz, & H. T. Ngo, Eds.). Intergovernmental Science-Policy Platform on Biodiversity and Ecosystem Services (IPBES).

IPBES. (2023). *Thematic Assessment Report on Invasive Alien Species and their Control of the Intergovernmental Science-Policy Platform on Biodiversity and Ecosystem Services*. (A. Pauchard, P. Stoett, H. E. Roy, & T. Renard Truong, Eds.; Version 4). 10.5281/ZENODO.7430682

James, K., & Bradshaw, K. (2020). Detecting plant species in the field with deep learning and drone technology. Methods in Ecology and Evolution, 11(11), 1509–1519. 10.1111/2041-210X.13473

Jiménez-Valverde, A., Peterson, A. T., Soberón, J., Overton, J. M., Aragón, P., & Lobo, J. M. (2011). Use of niche models in invasive species risk assessments. Biological Invasions, 13(12), 2785–2797. 10.1007/s10530-011-9963-4

Jodoin, Y., Lavoie, C., Villeneuve, P., Theriault, M., Beaulieu, J., & Belzile, F. (2008). Highways as corridors and habitats for the invasive common reed *Phragmites australis* in Quebec, Canada. Journal of Applied Ecology, 45(2), 459–466. 10.1111/j.1365-2664.2007.01362.x

Karathanos, S., Rivard, N., Brisson, J., & Lavoie, C. (2015). Limiter l’invasion du roseau commun sur des terres en friche. Bulletin de conservation, 4.

Katal, N., Rzanny, M., Mäder, P., & Wäldchen, J. (2022). Deep Learning in Plant Phenological Research: A Systematic Literature Review. Frontiers in Plant Science, 13, 805738. 10.3389/fpls.2022.805738

Kattenborn, T., Eichel, J., & Fassnacht, F. E. (2019). Convolutional Neural Networks enable efficient, accurate and fine-grained segmentation of plant species and communities from high-resolution UAV imagery. Scientific Reports, 9(1), 17656. 10.1038/s41598-019-53797-9

Kattenborn, T., Leitloff, J., Schiefer, F., & Hinz, S. (2021). Review on Convolutional Neural Networks (CNN) in vegetation remote sensing. ISPRS Journal of Photogrammetry and Remote Sensing, 173, 24–49. 10.1016/j.isprsjprs.2020.12.010

Kattenborn, T., Lopatin, J., Förster, M., Braun, A. C., & Fassnacht, F. E. (2019). UAV data as alternative to field sampling to map woody invasive species based on combined Sentinel-1 and Sentinel-2 data. Remote Sensing of Environment, 227, 61–73. 10.1016/j.rse.2019.03.025

Kettenring, K. M., & Adams, C. R. (2011). Lessons learned from invasive plant control experiments: A systematic review and meta-analysis: Invasive plant control experiments. Journal of Applied Ecology, 48(4), 970–979. 10.1111/j.1365-2664.2011.01979.x

Laliberté, E., Cogliastro, A., & Bouchard, A. (2006). Projet pilote de restauration de paysages forestiers au parc national des Îles-de-Boucherville (p. 57). Institut de recherche en biologie végétale.

Lavergne, S., & Molofsky, J. (2004). Reed Canary Grass (*Phalaris arundinacea*) as a Biological Model in the Study of Plant Invasions. Critical Reviews in Plant Sciences, 23(5), 415–429. 10.1080/07352680490505934

Lavoie, C. (2007). Le roseau commun au Québec: Enquête sur une invasion. Le Naturaliste Canadien, 131(2), 5–9.

Lavoie, C., Ayotte, G., & Groeneveld, E. (2019). *50 plantes envahissantes: Protéger la nature et l’agriculture*.

Lavoie, C., Guay, G., & Joerin, F. (2014). Une liste des plantes vasculaires exotiques nuisibles du Québec: Nouvelle approche pour la sélection des espèces et l’aide à la décision. Écoscience, 21(2), 133–156. 10.2980/21-2-3703

Le groupe Phragmites. (2012). Le roseau envahisseur: La dynamique, l’impact et le contrôle d’une invasion d’envergure. Le Naturaliste canadien, 136(3), 33–39. 10.7202/1009238ar

LeCun, Y., Bengio, Y., & Hinton, G. (2015). Deep learning. Nature, 521(7553), 436–444. 10.1038/nature14539

Leduc, M.-B., & Knudby, A. (2018). Mapping Wild Leek through the Forest Canopy Using a UAV. Remote Sensing, 10(2), 70. 10.3390/rs10010070

Lehan, N. E., Murphy, J. R., Thorburn, L. P., & Bradley, B. A. (2013). Accidental introductions are an important source of invasive plants in the continental United States. American Journal of Botany, 100, 8.

Lelong, B., Lavoie, C., Jodoin, Y., & Belzile, F. (2007). Expansion pathways of the exotic common reed (Phragmites australis): A historical and genetic analysis. Diversity and Distributions, 13(4), 430–437. 10.1111/j.1472-4642.2007.00351.x

Levine, J. M., Adler, P. B., & Yelenik, S. G. (2004). A meta-analysis of biotic resistance to exotic plant invasions: Biotic resistance to plant invasion. Ecology Letters, 7(10), 975– 989. 10.1111/j.1461-0248.2004.00657.x

Levine, J. M., & D’Antonio, C. M. (1999). Elton Revisited: A Review of Evidence Linking Diversity and Invasibility. Oikos, 87(1), 15. 10.2307/3546992

Levine, J. M., Vilà, M., Antonio, C. M. D., Dukes, J. S., Grigulis, K., & Lavorel, S. (2003). Mechanisms underlying the impacts of exotic plant invasions. Proceedings of the Royal Society of London. Series B: Biological Sciences, 270(1517), 775–781. 10.1098/rspb.2003.2327

Lisein, J., Michez, A., Claessens, H., & Lejeune, P. (2015). Discrimination of Deciduous Tree Species from Time Series of Unmanned Aerial System Imagery. PLOS ONE, 10(11), e0141006. 10.1371/journal.pone.0141006

Lopatin, J., Dolos, K., Kattenborn, T., & Fassnacht, F. E. (2019). How canopy shadow affects invasive plant species classification in high spatial resolution remote sensing. Remote Sensing in Ecology and Conservation, 5(4), 302–317. 10.1002/rse2.109

Maggiori, E., Tarabalka, Y., Charpiat, G., & Alliez, P. (2017). Convolutional Neural Networks for Large-Scale Remote-Sensing Image Classification. IEEE Transactions on Geoscience and Remote Sensing, 55(2), 645–657. 10.1109/TGRS.2016.2612821

Maheu-Giroux, M., & Blois, S. D. (2005). Mapping the invasive species Phragmites australis in linear wetland corridors. Aquatic Botany, 83(4), 310–320. 10.1016/j.aquabot.2005.07.002

Mal, T. K., & Narine, L. (2004). The biology of Canadian weeds. 129. Phragmites australis (Cav.) Trin. Ex Steud. Canadian Journal of Plant Science, 84(1), 365–396. 10.4141/P01-172

Matese, A., Toscano, P., Di Gennaro, S., Genesio, L., Vaccari, F., Primicerio, J., Belli, C., Zaldei, A., Bianconi, R., & Gioli, B. (2015). Intercomparison of UAV, Aircraft and Satellite Remote Sensing Platforms for Precision Viticulture. Remote Sensing, 7(3), 2971–2990. 10.3390/rs70302971

Mehta, S. V., Haight, R. G., Homans, F. R., Polasky, S., & Venette, R. C. (2007). Optimal detection and control strategies for invasive species management. Ecological Economics, 61(2–3), 237–245. 10.1016/j.ecolecon.2006.10.024

Milas, A. S., Arend, K., Mayer, C., Simonson, M. A., & Mackey, S. (2017). Different colours of shadows: Classification of UAV images. International Journal of Remote Sensing, 38(8–10), 3084–3100. 10.1080/01431161.2016.1274449

Minchin, D., & Gollasch, S. (2003). Fouling and Ships’ Hulls: How Changing Circumstances and Spawning Events may Result in the Spread of Exotic Species. Biofouling, 19(sup1), 111–122. 10.1080/0892701021000057891

Ministère de l’Environnement et de la Lutte contre les changements climatiques. (2022). *Utilisation des pesticides en milieu aquatique—Guide d’apprentissage*. 67.

Mohler, R. L., & Morse, J. M. (2022). Using UAV imagery to map invasive Phragmites australis on the Crow Island State Game Area, Michigan, USA. Wetlands Ecology and Management, 30(6), 1213–1229. 10.1007/s11273-022-09890-4

Morrison, L. W. (2016). Observer error in vegetation surveys: A review. Journal of Plant Ecology, 9(4), 367–379. 10.1093/jpe/rtv077

MRC de Kamouraska. (2014). *Plantes exotiques envahissantes. Lutter contre l’invasion*. http://www.mrckamouraska.com/documentation/Presentationpee.pdf

Müllerová, J., Bartaloš, T., Brůna, J., Dvořák, P., & Vítková, M. (2017). Unmanned aircraft in nature conservation: An example from plant invasions. International Journal of Remote Sensing, 38(8–10), 2177–2198. 10.1080/01431161.2016.1275059

Müllerová, J., Brůna, J., Bartaloš, T., Dvořák, P., Vítková, M., & Pyšek, P. (2017). Timing Is Important: Unmanned Aircraft vs. Satellite Imagery in Plant Invasion Monitoring. Frontiers in Plant Science, 8, 887. 10.3389/fpls.2017.00887

Mumby, P. J., Green, E. P., Edwards, A. J., & Clark, C. D. (1999). The cost-effectiveness of remote sensing for tropical coastal resources assessment and management. Journal of Environmental Management, 55(3), 157–166. 10.1006/jema.1998.0255

Nantel, P., Claudi, R., & Muckle-Jeffs, E. (2002). *Envahisseurs exotiques des eaux, milieux humides et forêts du Canada*. Service canadien des forêts, Direction générale des sciences.

Nezami, S., Khoramshahi, E., Nevalainen, O., Pölönen, I., & Honkavaara, E. (2020). Tree Species Classification of Drone Hyperspectral and RGB Imagery with Deep Learning Convolutional Neural Networks. Remote Sensing, 12(7), 1070. 10.3390/rs12071070

Over, J.-S. R., Ritchie, A. C., Kranenburg, C. J., Brown, J. A., Buscombe, D. D., Noble, T., Sherwood, C. R., Warrick, J. A., & Wernette, P. A. (2021). Processing coastal imagery with Agisoft Metashape Professional Edition, version 1.6—Structure from motion workflow documentation. In Open-File Report (2021–1039). U.S. Geological Survey. 10.3133/ofr20211039

Packer, J. G., Meyerson, L. A., Skálová, H., Pyšek, P., & Kueffer, C. (2017). Biological Flora of the British Isles: *Phragmites australis*. Journal of Ecology, 105(4), 1123–1162. 10.1111/1365-2745.12797

Park, M. G., & Blossey, B. (2008). Importance of plant traits and herbivory for invasiveness of *Phragmites australis* (Poaceae). American Journal of Botany, 95(12), 1557–1568. 10.3732/ajb.0800023

Payne, J. L., Bush, A. M., Heim, N. A., Knope, M. L., & McCauley, D. J. (2016). Ecological selectivity of the emerging mass extinction in the oceans. Science, 353(6305), 1284–1286. 10.1126/science.aaf2416

Pejchar, L., & Mooney, H. A. (2009). Invasive species, ecosystem services and human well-being. Trends in Ecology & Evolution, 24(9), 497–504. 10.1016/j.tree.2009.03.016

Pellerin, S., Duquesne, T., Omelczuk Walter, C., & Pasquet, S. (2016). La richesse floristique des friches du Parc national de Frontenac. Le Naturaliste canadien, 141(1), 15–23. 10.7202/1037933ar

Peng, S., Kinlock, N. L., Gurevitch, J., & Peng, S. (2019). Correlation of native and exotic species richness: A global meta-analysis finds no invasion paradox across scales. Ecology, 100(1). 10.1002/ecy.2552

Pengra, B. W., Johnston, C. A., & Loveland, T. R. (2007). Mapping an invasive plant, Phragmites australis, in coastal wetlands using the EO-1 Hyperion hyperspectral sensor. Remote Sensing of Environment, 108(1), 74–81. 10.1016/j.rse.2006.11.002

Pimentel, D., Zuniga, R., & Morrison, D. (2005). Update on the environmental and economic costs associated with alien-invasive species in the United States. Ecological Economics, 52(3), 273–288. 10.1016/j.ecolecon.2004.10.002

Potts, B. M., Barbour, R. C., Hingston, A. B., & Vaillancourt, R. E. (2003). Genetic pollution of native eucalypt gene pools—Identifying the risks. Australian Journal of Botany, 51(1), 1. 10.1071/BT02035

Prince, S. J. D. (2012). Computer Vision: Models, Learning, and Inference. Cambridge University Press.

Pyšek, P., Hulme, P. E., Simberloff, D., Bacher, S., Blackburn, T. M., Carlton, J. T., Dawson, W., Essl, F., Foxcroft, L. C., Genovesi, P., Jeschke, J. M., Kühn, I., Liebhold, A. M., Mandrak, N. E., Meyerson, L. A., Pauchard, A., Pergl, J., Roy, H. E., Seebens, H., … Richardson, D. M. (2020). Scientists’ warning on invasive alien species. Biological Reviews, 95(6), 1511–1534. 10.1111/brv.12627

Reaser, J. K., Burgiel, S. W., Kirkey, J., Brantley, K. A., Veatch, S. D., & Burgos-Rodríguez, J. (2020). The early detection of and rapid response (EDRR) to invasive species: A conceptual framework and federal capacities assessment. Biological Invasions, 22(1), 1–19. 10.1007/s10530-019-02156-w

Reckling, W., Mitasova, H., Wegmann, K., Kauffman, G., & Reid, R. (2021). Efficient Drone-Based Rare Plant Monitoring Using a Species Distribution Model and AI-Based Object Detection. Drones, 5(4), 110. 10.3390/drones5040110

Reichard, S. H., & White, P. (2001). Horticulture as a Pathway of Invasive Plant Introductions in the United States. BioScience, 51(2), 103. 10.1641/0006-3568(2001)051 [0103:HAAPOI]2.0.CO;2

Richter, M. L., Byttner, W., Krumnack, U., Wiedenroth, A., Schallner, L., & Shenk, J. (2021). (Input) Size Matters for CNN Classifiers. In I. Farkaš, P. Masulli, S. Otte, & S. Wermter (Eds.), Artificial Neural Networks and Machine Learning – ICANN 2021 (Vol. 12892, pp. 133–144). Springer International Publishing. 10.1007/978-3-030-86340-1_11

Rinella, M. J., Maxwell, B. D., Fay, P. K., Weaver, T., & Sheley, R. L. (2009). Control effort exacerbates invasive-species problem. Ecological Applications, 19(1), 155–162. 10.1890/07-1482.1

Ross, M. (1990). *Étude sur l’évolution de boisé Grosbois au Parc des îles-de-Boucherville*. Parcs de la Montérégie, Service de la gestion des ressources naturelles.

Safonova, A., Tabik, S., Alcaraz-Segura, D., Rubtsov, A., Maglinets, Y., & Herrera, F. (2019). Detection of Fir Trees (Abies sibirica) Damaged by the Bark Beetle in Unmanned Aerial Vehicle Images with Deep Learning. Remote Sensing, 11(6), 643. 10.3390/rs11060643

Samiappan, S., Turnage, G., Hathcock, L., Casagrande, L., Stinson, P., & Moorhead, R. (2017). Using unmanned aerial vehicles for high-resolution remote sensing to map invasive *Phragmites australis* in coastal wetlands. International Journal of Remote Sensing, 38(8–10), 2199–2217. 10.1080/01431161.2016.1239288

Schiefer, F., Kattenborn, T., Frick, A., Frey, J., Schall, P., Koch, B., & Schmidtlein, S. (2020). Mapping forest tree species in high resolution UAV-based RGB-imagery by means of convolutional neural networks. ISPRS Journal of Photogrammetry and Remote Sensing, 170, 205–215. 10.1016/j.isprsjprs.2020.10.015

Seebens, H., Blackburn, T. M., Dyer, E. E., Genovesi, P., Hulme, P. E., Jeschke, J. M., Pagad, S., Pyšek, P., Winter, M., Arianoutsou, M., Bacher, S., Blasius, B., Brundu, G., Capinha, C., Celesti-Grapow, L., Dawson, W., Dullinger, S., Fuentes, N., Jäger, H., … Essl, F. (2017). No saturation in the accumulation of alien species worldwide. Nature Communications, 8(1), 14435. 10.1038/ncomms14435

Sépaq. (2021). *Bulletin de conservation*. https://www.sepaq.com/resources/docs/pq/pq_bulletin_2021.pdf

Sépaq. (2022). *Plan de conservation 2022-2027 – Parc national des Îles-de-Boucherville et parc national du Mont-Saint-Bruno*. Sépaq. https://imagescloud.s3.amazonaws.com/documents/parcs-nationaux/conservation/plans-conservation/bou_msb_plan_conservation_2022-2027.pdf

Shea, K., & Chesson, P. (2002). Community ecology theory as a framework for biological invasions. Trends in Ecology & Evolution, 17(4), 170–176. 10.1016/S0169-5347(02)02495-3

Simberloff, D., Martin, J.-L., Genovesi, P., Maris, V., Wardle, D. A., Aronson, J., Courchamp, F., Galil, B., García-Berthou, E., Pascal, M., Pyšek, P., Sousa, R., Tabacchi, E., & Vilà, M. (2013). Impacts of biological invasions: What’s what and the way forward. Trends in Ecology & Evolution, 28(1), 58–66. 10.1016/j.tree.2012.07.013

Sokolova, M., Japkowicz, N., & Szpakowicz, S. (2006). Beyond Accuracy, F-Score and ROC: A Family of Discriminant Measures for Performance Evaluation. In A. Sattar & B. Kang (Eds.), AI 2006: Advances in Artificial Intelligence (Vol. 4304, pp. 1015–1021). Springer Berlin Heidelberg. 10.1007/11941439_114

Sun, Z., Wang, X., Wang, Z., Yang, L., Xie, Y., & Huang, Y. (2021). UAVs as remote sensing platforms in plant ecology: Review of applications and challenges. Journal of Plant Ecology, 14(6), 1003–1023. 10.1093/jpe/rtab089

Takaya, K., Sasaki, Y., & Ise, T. (2022). Automatic detection of alien plant species in action camera images using the chopped picture method and the potential of citizen science. Breeding Science, 21062. 10.1270/jsbbs.21062

Thomas, C. D., & Palmer, G. (2015). Non-native plants add to the British flora without negative consequences for native diversity. Proceedings of the National Academy of Sciences, 112(14), 4387–4392. 10.1073/pnas.1423995112

Tomasetto, F., Duncan, R. P., & Hulme, P. E. (2019). Resolving the invasion paradox: Pervasive scale and study dependence in the native-alien species richness relationship. Ecology Letters, 22(6), 1038–1046. 10.1111/ele.13261

Tougas-Tellier, M.-A. (2013). Impact des changements climatiques sur l’expansion du roseau envahisseur dans le fleuve Saint-Laurent. Université Laval.

Vilà, M., Espinar, J. L., Hejda, M., Hulme, P. E., Jarošík, V., Maron, J. L., Pergl, J., Schaffner, U., Sun, Y., & Pyšek, P. (2011). Ecological impacts of invasive alien plants: A meta-analysis of their effects on species, communities and ecosystems: Ecological impacts of invasive alien plants. Ecology Letters, 14(7), 702–708. 10.1111/j.1461-0248.2011.01628.x

Vymazal, J., & Krőpfelová, L. (2005). Growth of Phragmites australis and Phalaris arundinacea in constructed wetlands for wastewater treatment in the Czech Republic. Ecological Engineering, 25(5), 606–621. 10.1016/j.ecoleng.2005.07.005

Wang, K., Zhang, D., Li, Y., Zhang, R., & Lin, L. (2017). Cost-Effective Active Learning for Deep Image Classification. IEEE Transactions on Circuits and Systems for Video Technology, 27(12), 2591–2600. 10.1109/TCSVT.2016.2589879

Wang, S., Guan, K., Zhang, C., Zhou, Q., Wang, S., Wu, X., Jiang, C., Peng, B., Mei, W., Li, K., Li, Z., Yang, Y., Zhou, W., Huang, Y., & Ma, Z. (2023). Cross-scale sensing of field-level crop residue cover: Integrating field photos, airborne hyperspectral imaging, and satellite data. Remote Sensing of Environment, 285, 113366. 10.1016/j.rse.2022.113366

Weil, G., Lensky, I., Resheff, Y., & Levin, N. (2017). Optimizing the Timing of Unmanned Aerial Vehicle Image Acquisition for Applied Mapping of Woody Vegetation Species Using Feature Selection. Remote Sensing, 9(11), 1130. 10.3390/rs9111130

Weisberg, P. J., Dilts, T. E., Greenberg, J. A., Johnson, K. N., Pai, H., Sladek, C., Kratt, C., Tyler, S. W., & Ready, A. (2021). Phenology-based classification of invasive annual grasses to the species level. Remote Sensing of Environment, 263, 112568. 10.1016/j.rse.2021.112568

Williamson, M., & Fitter, A. (1996). The Varying Success of Invaders. Ecology, 77(6), 1661– 1666. 10.2307/2265769

Wolkovich, E. M., & Cleland, E. E. (2011). The phenology of plant invasions: A community ecology perspective. Frontiers in Ecology and the Environment, 9(5), 287–294. 10.1890/100033

Xie, Y., Sha, Z., & Yu, M. (2008). Remote sensing imagery in vegetation mapping: A review. Journal of Plant Ecology, 1(1), 9–23. 10.1093/jpe/rtm005

Zhang, L., Zhang, L., & Du, B. (2016). Deep Learning for Remote Sensing Data: A Technical Tutorial on the State of the Art. IEEE Geoscience and Remote Sensing Magazine, 4(2), 22–40. 10.1109/MGRS.2016.2540798

