## Supplementary material for "Early Detection of an Invasive Alien Plant (*Phragmites australis*) Using Unoccupied Aerial Vehicles and Artificial Intelligence"

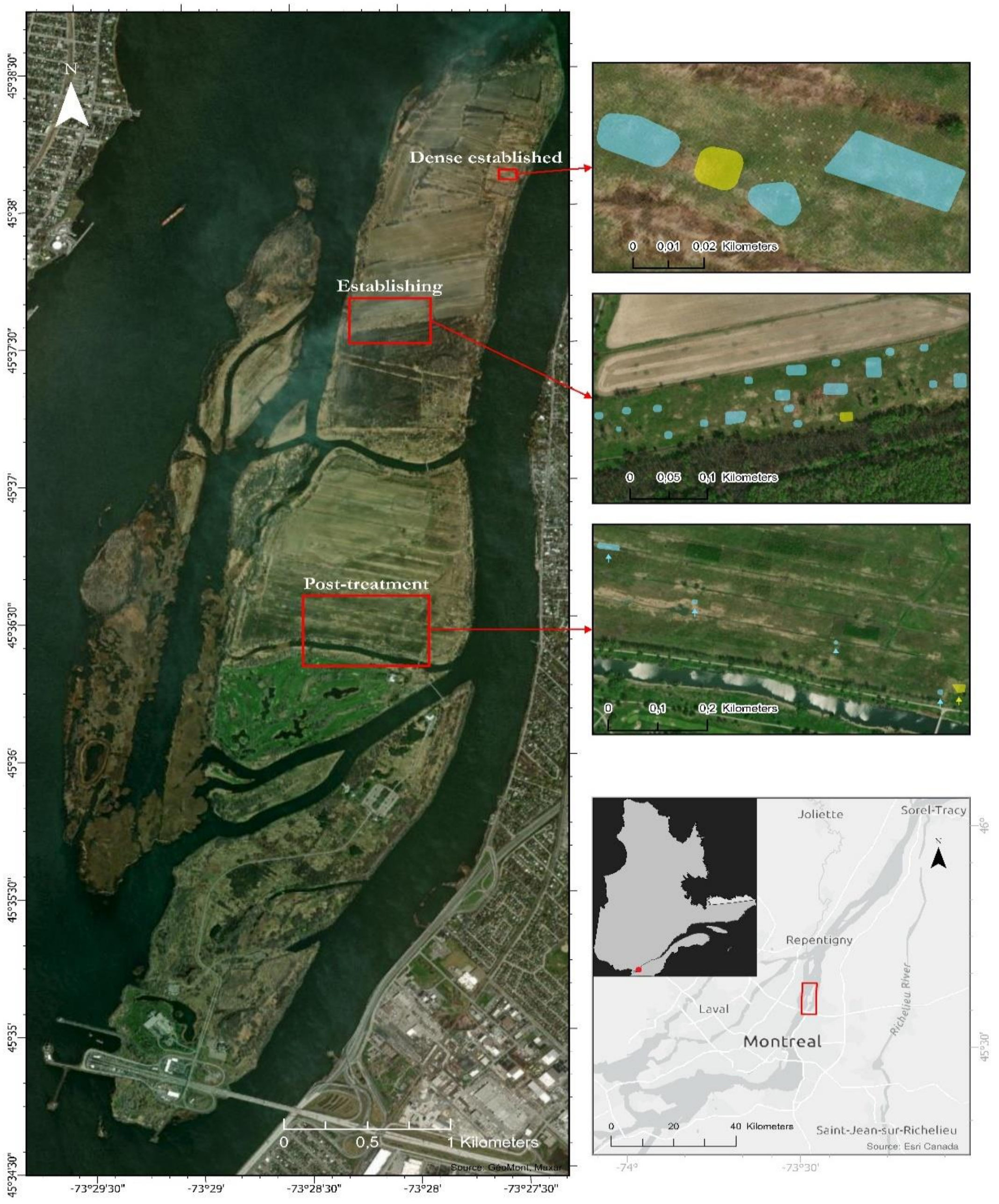


**Supplementary Figure S1** Map of the Îles-de-Boucherville, Québec, Canada with the location of the plots

The polygons in blue represent the plots used for models training and validation while the polygons in yellow represent the data used to test the final model with unseen data. Projection: WGS84 UTM Zone 18N (EPSG: 32618). The subset at bottom right indicates the position of the park in relation to the city of Montréal (red rectangle) and within the province of Quebec (red dot).

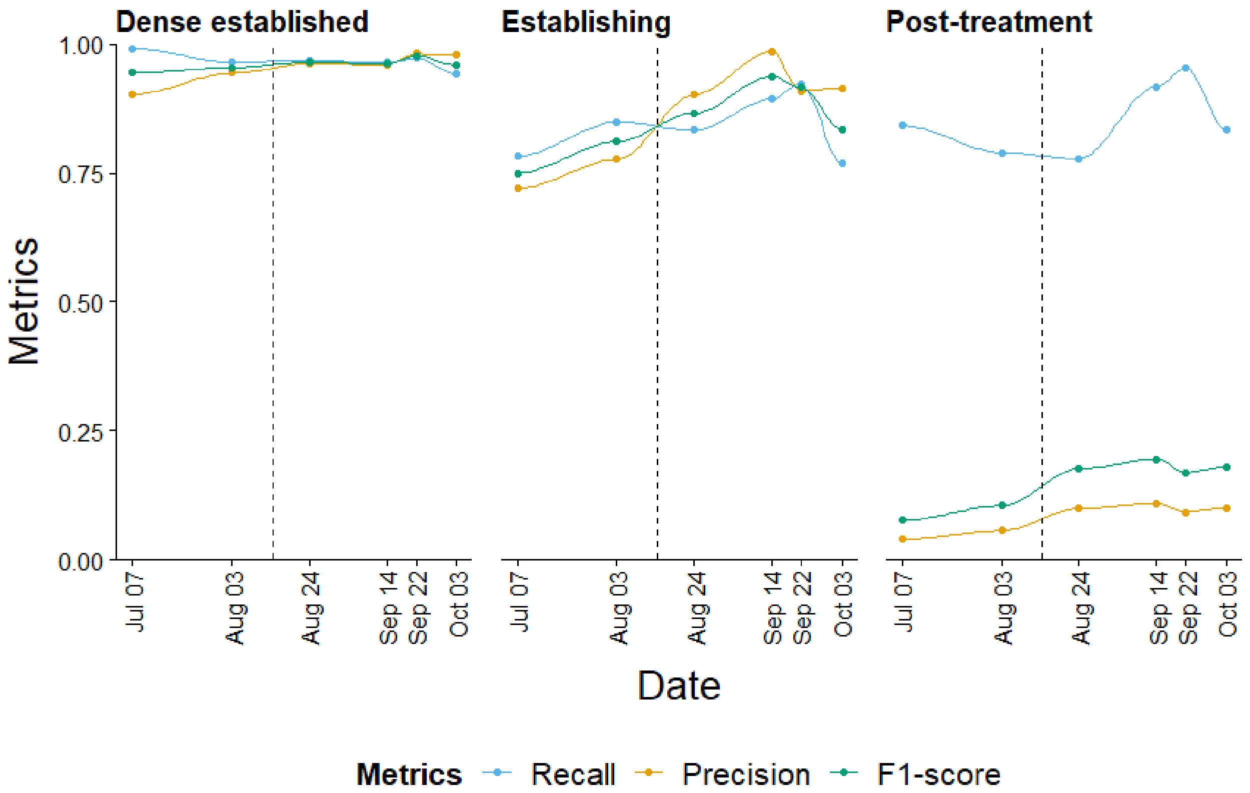


**Supplementary Figure S2** Evolution of recall, precision and F1-score for each zone at each date

The dotted vertical line indicates the approximate moment (between Aug. 3 and Aug. 24) at which the common reed inflorescence emerges and becomes visible in the imagery. There is a general increase in the F1-score metric throughout the growing season and a peak is reached in September, followed by a decrease for the last date.

**Supplementary Table S1** Performance measures for each model parameter tested ahead of final training

Tile size, number of epochs, batch size, loss function used, optimizer used and model architecture used with two different pre-training datasets were the parameters tested. The default values for each parameter while testing were 256 x 256 pixels, 20 epochs, 48 tiles, Focal Loss, Adam, and VGG19_BN with ImageNet respectively. Model architecture trained with ImageNet (Deng et al., 2009) are all implanted by default in fastai library while those trained with PlantNet – 300k are from Garcin et al. (2021). All results for the tested parameters are from predictions on the validation dataset. The highest F1-score for each parameter is shown in bold and corresponds to the selected value.

| Parameter | Values | | Recall | Precision | F1-Score |
|---|---|---|---|---|---|
| **Tile size** | 128 x 128 | | 0.953 | 0.933 | 0.943 |
| | **256 x 256** | | **0.955** | **0.959** | **0.957** |
| | 512 x 512 | | 0.908 | 0.979 | 0.942 |
| **Epoch** | 10 | | 0.961 | 0.963 | 0.962 |
| | 15 | | 0.971 | 0.951 | 0.961 |
| | **20** | | **0.971** | **0.956** | **0.964** |
| | 25 | | 0.981 | 0.939 | 0.960 |
| | 30 | | 0.980 | 0.941 | 0.960 |
| **Batch size** | 4 | | 0.911 | 0.969 | 0.939 |
| | 8 | | 0.954 | 0.948 | 0.951 |
| | 24 | | 0.971 | 0.931 | 0.951 |
| | 32 | | 0.975 | 0.925 | 0.949 |
| | 48 | | 0.972 | 0.932 | 0.951 |
| | **72** | | **0.971** | **0.956** | **0.964** |
| | 96 | | 0.975 | 0.915 | 0.944 |
| **Loss function** | Cross Entropy | | 0.974 | 0.936 | 0.955 |
| | **Focal Loss** | | **0.971** | **0.956** | **0.964** |
| **Optimizer** | **Adam** | | **0.971** | **0.956** | **0.964** |
| | Lion | | 0.939 | 0.945 | 0.942 |
| **Model architecture** | AlexNet [1] | ImageNet | 0.940 | 0.864 | 0.900 |
| | DenseNet121 [2] | ImageNet | 0.969 | 0.921 | 0.944 |
| | DenseNet161 [2] | ImageNet | 0.974 | 0.929 | 0.951 |
| | DenseNet169 [2] | ImageNet | 0.971 | 0.920 | 0.945 |
| | DenseNet201 [2] | ImageNet | 0.967 | 0.942 | 0.954 |
| | ResNet34 [3] | ImageNet | 0.971 | 0.902 | 0.936 |
| | ResNet50 [3] | ImageNet | 0.975 | 0.907 | 0.940 |
| | ResNet101 [3] | ImageNet | 0.976 | 0.910 | 0.942 |
| | ResNet152 [3] | ImageNet | 0.974 | 0.916 | 0.944 |

| | | | | |
|---|---|---|---|---|
| SqueezeNet1_0 [4] | ImageNet | 0.957 | 0.902 | 0.929 |
| SqueezeNet1_1 [4] | ImageNet | 0.950 | 0.889 | 0.918 |
| VGG11 [5] | ImageNet | 0.962 | 0.891 | 0.925 |
| VGG16 [5] | ImageNet | 0.961 | 0.926 | 0.943 |
| VGG19 [5] | ImageNet | 0.963 | 0.923 | 0.942 |
| VGG16_BN [5] | ImageNet | 0.970 | 0.930 | 0.950 |
| **VGG19_BN** [5] | **ImageNet** | **0.971** | **0.938** | **0.955** |
| AlexNet [1] | PlantNet - 300K | 0.914 | 0.853 | 0.883 |
| ResNet18 [3] | PlantNet - 300K | 0.948 | 0.900 | 0.923 |
| ResNet34 [3] | PlantNet - 300K | 0.955 | 0.911 | 0.932 |
| ResNet50 [3] | PlantNet - 300K | 0.961 | 0.930 | 0.945 |
| ResNet101 [3] | PlantNet - 300K | 0.963 | 0.919 | 0.941 |
| ResNet152 [3] | PlantNet - 300K | 0.964 | 0.926 | 0.945 |
| VGG11[5] | PlantNet - 300K | 0.950 | 0.918 | 0.934 |
| WideResNet50_2 [6] | PlantNet - 300K | 0.960 | 0.937 | 0.949 |
| WideResNet101_2 [6] | PlantNet - 300K | 0.963 | 0.929 | 0.946 |


[1] Krizhevsky, A. (2014). One weird trick for parallelizing convolutional neural networks (arXiv:1404.5997). arXiv. http://arxiv.org/abs/1404.5997

[2] Huang, G., Liu, Z., van der Maaten, L., & Weinberger, K. Q. (2018). *Densely Connected Convolutional Networks* (arXiv:1608.06993). arXiv. https://doi.org/10.48550/arXiv.1608.06993

[3] He, K., Zhang, X., Ren, S., & Sun, J. (2015). *Deep Residual Learning for Image Recognition* (arXiv:1512.03385). arXiv. http://arxiv.org/abs/1512.03385

[4] Iandola, F. N., Han, S., Moskewicz, M. W., Ashraf, K., Dally, W. J., & Keutzer, K. (2016). *SqueezeNet: AlexNet-level accuracy with 50x fewer parameters and <0.5MB model size* (arXiv:1602.07360). arXiv. https://doi.org/10.48550/arXiv.1602.07360

[5] Simonyan, K., & Zisserman, A. (2015). *Very Deep Convolutional Networks for Large-Scale Image Recognition* (arXiv:1409.1556). arXiv. https://doi.org/10.48550/arXiv.1409.1556

[6] Zagoruyko, S., & Komodakis, N. (2017). *Wide Residual Networks* (arXiv:1605.07146). arXiv. https://doi.org/10.48550/arXiv.1605.07146

**Supplementary Table S2** Sky and wind conditions for each zone at each date

The meteorological conditions were not always uniform throughout the entire duration of a mission. For instance, it was possible to encounter both cloudy and clear skies during the same drone flight. These values represent the average conditions for each zone on each date. Wind conditions were determined from the values transmitted by the drone and converted to the Beaufort wind force scale.

| Zone | Date | Sky conditions | Wind force (Beaufort scale) |
|---|---|---|---|
| **Dense established** | **July 07** | Partly cloudy | 2 |
| | **Aug. 03** | Clear sky | 3 |
| | **Aug. 24** | Clear sky | 1 |
| | **Sept. 14** | Partly cloudy | 5 |
| | **Sept. 22** | Fully cloudy | 3 |
| | **Oct. 03** | Clear sky | 2 |
| **Establishing** | **July 07** | Clear sky | 2 |
| | **Aug. 03** | Partly cloudy | 4 |
| | **Aug. 24** | Clear sky | 1 |
| | **Sept. 14** | Clear sky | 5 |
| | **Sept. 22** | Fully cloudy | 5 |
| | **Oct. 03** | Clear sky | 2 |
| **Post-treatment** | **July 07** | Clear sky | 2 |
| | **Aug. 03** | Partly cloudy | 2 |
| | **Aug. 24** | Partly cloudy | 2 |
| | **Sept. 14** | Partly cloudy | 5 |
| | **Sept. 22** | Fully cloudy | 3 |
| | **Oct. 03** | Clear sky | 2 |
